# Local climate and vernalization requirements explain the latitudinal patterns of flowering initiation in the crop wild relative *Linum bienne*

**DOI:** 10.1101/2022.01.02.474722

**Authors:** Beatrice Landoni, Pilar Suárez-Montes, Rico H. F. Habeahan, Adrian C. Brennan, Rocío Pérez-Barrales

## Abstract

**Background and Aims:** Days to flowering initiation in species with large geographic distribution often correlate with latitude. Latitude reflects climatic gradients, but it is unclear if large-scale differentiation in flowering results from adaptation to local climate, and whether adaptation to local climate could constrain shifts in distribution and colonization of new environments.

**Methods:** In its Western range in Europe, *L. bienne* populations were surveyed to describe latitudinal patterns of flowering initiation and determine its correlation with the local climate of populations. This was measured under standardized greenhouse conditions, with a vernalization experiment to learn if chilling advances flowering, and with a reciprocal transplant experiment at three sites along the latitudinal gradient, recording flowering at the central site and plant survival in all sites. Also, genetic differentiation of populations along the latitudinal range was studied using microsatellite markers.

**Key Results:** Flowering initiation varied with latitude, with southern populations flowering earlier than northern populations. Latitude also predicted population response to vernalization, with chilling inducing a greater advance of flowering initiation in northern than southern populations. In general, plant survival in the reciprocal transplant experiment decreased with the geographic distance of populations to the experimental site and, at the central site, flowering initiation varied with latitude of origin. However, across experiments, the local climate of populations better predicted the differentiation in flowering initiation and vernalization response than latitude of origin. Finally, the microsatellite data revealed genetic differentiation of populations forming two groups that agree with a Mediterranean and Atlantic lineage.

**Conclusions:** The consistent result across experiments of a latitudinal cline in flowering initiation and in the vernalization response suggests that flowering is under genetic regulation and yet dependent on particular environmental and climatic cues at local scale. However, the genetic differentiation suggests that past population history might influenced the flowering initiation patterns detected.

## Introduction

Flowering initiation in temperate species with wide geographic distribution varies with latitude (Boudry *et al*. 2002; Olsson and Ågren 2002; Stinchcombe *et al*. 2004; Debieu *et al*. 2013; Richardson *et al*. 2016). This variation is partially explained by the fact that latitude reflects environmental gradients including photoperiod, temperature, and precipitation that often exert selection on flowering phenology and other life history traits (Keller *et al*. 2009; Pau *et al*. 2011; Colautti and Barrett 2013; de Frenne *et al*. 2013; Preite *et al*. 2015; Burgarella *et al*. 2016; Muir and Angert 2017). Flowering initiation at finer spatial scales correlates with altitudinal gradients or topological heterogeneity that also modify the local climate (Morente-López *et al*.; Franks *et al*. 2007; Mendez-Vigo *et al*. 2011; Halbritter *et al*. 2018; Lampei *et al*. 2019). Since the time of flowering affects the start of following reproductive events, and therefore plant fitness, the spatial variation in flowering initiation is seen as evidence of adaptation to environmental variation at multiple geographical scales (Morente-López *et al*.; Endler 1977; Boudry *et al*. 2002; Stinchcombe *et al*. 2004; Mendez-Vigo *et al*. 2011; Burgarella *et al*. 2016). The adaptive nature of flowering initiation has been tested with the use of reciprocal transplant experiments, often revealing that local adaptation is an important mechanism creating clinal variation in flowering (Ågren and Schemske 2012; Colautti and Barrett 2013; Lowry *et al*. 2019).

Flowering initiation is a trait under the tight regulation of a complex network of genes that respond to endogenous and environmental cues. In this way, plants can synchronise reproduction with the optimal local growing conditions to maximise fitness across a range of environments (Amasino and Michaels 2010; Salomé *et al*. 2011; Andrés and Coupland 2012; Blackman 2017; Bouché *et al*. 2017; Zan and Carlborg 2019). Among the different flowering pathways, vernalization and photoperiod are perhaps the most relevant in temperate species because variation in temperature and daylength are reliable cues of seasonal transitions (Amasino and Michaels 2010; Andrés and Coupland 2012; Blackman 2017; Whittaker and Dean 2017; Friedman 2020). This is evidenced by the spatial distribution of allelic variation of genes involved in the vernalization and photoperiod pathways along environmental clines (Stinchcombe *et al*. 2004, 2005; Samis *et al*. 2008; Keller *et al*. 2011; Mendez-Vigo *et al*. 2011; Burgarella *et al*. 2016). In addition to genetic regulation, environmental variation can induce adaptive plastic responses in flowering initiation and in other phenological traits (Sultan 2000; Ghalambor *et al*. 2007; Nicotra *et al*. 2010). These adaptive plastic responses often vary clinally (Stinchcombe *et al*. 2005; Mendez-Vigo *et al*. 2011; Lewandowska-Sabat *et al*. 2012; Toftegaard *et al*. 2016; Prevéy *et al*. 2017; Thibault *et al*. 2020). For example, responses of flowering initiation to vernalization have been associated with adaptation to different climates in *A. thaliana*, resulting in differential timing of flowering and other life-cycle events across its range (Exposito-alonso; Whittaker and Dean 2017), a pattern also detected in other herbaceous species in the wild (Boudry *et al*. 2002; Quilot-Turion *et al*. 2013). In sea beet, a latitudinal cline in vernalization requirement for flowering correlates with temperature and disturbance, in turn producing both fast-cycling annual and perennial life-histories across its native range (Boudry *et al*. 2002; Hautekèete *et al*. 2002). Hence, population variation in response to vernalization in temperate species with a large geographic range often corresponds with variation in the life cycle and life-history traits (Exposito-alonso; Boudry *et al*. 2002; Gremer *et al*. 2020; Friedman 2020).

With many wild plant species already responding to climate change (Sheth and Angert 2018; Midolo and Wellstein 2020; Anderson and Wadgymar 2020), learning the causes of differentiation in flowering initiation is critical to predict how species persist in and respond to a changing environment (Valladares *et al*. 2014; Ehrlén and Valdés 2020). Variation in flowering initiation can be explained by the correlations with other traits (Colautti *et al*. 2010; Haselhorst *et al*. 2011; Ehrlén 2015; Auge *et al*. 2019), or by neutral processes (Leimu and Fischer 2008; Keller *et al*. 2009; Chen *et al*. 2012). Standing genetic variation and phenotypic plasticity of flowering traits allow fast responses to abrupt changes in climate (Franks *et al*. 2007; van Dijk 2009; van Dijk and Hautekèete 2014). In temperate species, the timing of phenological phases, especially the shift from vegetative growth to initiation of flowering, is often dependent on the extent of vernalization and photoperiod experienced, the developmental stage of plants, and dependencies between traits (Boudry *et al*. 2002; Stinchcombe *et al*. 2004, 2005; Quilot-Turion *et al*. 2013; Rubin and Friedman 2018; Thibault *et al*. 2020; Gremer *et al*. 2020). With rising temperatures, how plant populations experience vernalization relative to photoperiod has changed, in turn shifting the initiation of flowering and the phenology of correlated traits with potential ecological consequences (Cook *et al*. 2012; van Dijk and Hautekèete 2014).

*Linum bienne* Mill., is an herbaceous species variously described as annual, winter annual, biennial, or perennial, and with a wide geographic range spanning the entire Mediterranean Basin and Western Europe, encompassing diverse environments (Diederichsen and Hammer 1995; Weiss and Zohary 2011; Uysal *et al*. 2012; Soto-Cerda *et al*. 2014). Natural variation in *L. bienne* remains poorly described but studies of Turkish populations and wider germplasm collections have found that geographic distance and altitude correlate with variation in flowering initiation and other traits, suggesting that this variation could be adaptive (Uysal *et al*. 2010, 2012; Soto-Cerda *et al*. 2014). *Linum bienne* is the wild progenitor of cultivated flax, *L. usitatissimum*, a crop grown across temperate regions of the world (Zohary and Hopf 2000; Weiss and Zohary 2011). Variation in responses to environmental conditions can also determine variation in flowering time of cultivated flax. Globally, associations between genomic regions and latitude, daylength, or mean daily temperature have been found within genetic clusters of cultivated flax (Gutaker *et al*. 2019; Sertse *et al*. 2019). The crop also shows variation in its response to vernalization and photoperiod which influences flowering in varieties grown at different latitudes in the USA (Fu 2012; Darapuneni *et al*. 2014). This is not uncommon in crops from other plant families where variation in vernalization requirement is fundamental to enable the cultivation of a crop under different environmental conditions (Saisho *et al*.; Abbo *et al*. 2002; Casao *et al*. 2011; Adhikari *et al*. 2012; Höft *et al*. 2018).

The present study quantifies the variation present in *L. bienne* for flowering initiation and its response to vernalization across a wide latitudinal and climatic range in Europe. The expectation is that flowering initiation, its response to vernalization, and consequent fitness co-vary with latitude, partly due to adaptation to the local climate that populations experience. To this end, *L. bienne* populations sampled along the western range of the species native range were exposed to various environments in experiments performed under glasshouse and field conditions (common garden, vernalization, and reciprocal transplant experiment). Six populations representative of the latitudinal range were also genotyped with microsatellite markers to determine population genetic structure against which to compare phenotypic trait variation. Moreover, *L. bienne* was compared to *L. usitatissimum* with the expectation that *L. bienne* displays more variation than its cultivated relative for flowering initiation and its response to vernalization. To this end, a collection of *L. usitatissimum* cultivars was included in the vernalization experiment.

## Material and Methods

### Study species and population collection

*Linum bienne* Mill. is an annual, biannual, or perennial herb, with a wide geographic distribution, ranging the entire Mediterranean Region, Atlantic Europe and the British Isles. It is a grassland species growing in dry and calcareous soils, from sea level up to 1200m. Plants present narrow and slender stems, with cymose inflorescence bearing pale blue flowers. Flowers are homostylous, self-compatible and experience autonomous self-pollination (upon flower opening, stigmas are receptive and dehisced anthers contact stigmas, B. Landoni and R. Perez-Barrales, personal observation), similar to cultivated flax (Williams *et al*. 1990). Fruits are small capsules bearing up to 10 seeds (Martínez-Labarga and Muñoz-Garmendia 2015). In this study, online databases (Anthos: http://www.anthos.es/, GBIF: https://www.gbif.org/, and BSBI: https://database.bsbi.org/) were used to plan the population survey in the western latitudinal range of the species distribution (Sicily, Spain, France, and England). Only for one population (Dor) bulked seed was obtained from Emorsgate Seeds, King’s Lynn, UK (https://wildseed.co.uk). A total of 34 populations were surveyed (10 to 42 individual plants per population; Fig. 1A and supplementary table 1 for details on geographic coordinates and elevation) between 2013 and 2017, although most populations were collected in 2016. Field samples were obtained by collecting fruits from a single individual plant (hereafter family) per patch, allowing at least 1 m distance between patches to avoid sampling the same maternal individual. At the time of collection, all families presented fruit capsules close to ripening. Collected families were placed in individual paper envelopes and stored in a cold room at 4°C with silica gel to absorb the excess of humidity. Seeds of 16 cultivated varieties originating from across western European and Canada were provided from Flaxland (Stroud, UK), Terre de Lin (Saint-Pierre-le-Viger, France), and Leibniz Institute of Plant Genetics and Crop Plant Research (Gatersleben, Germany). Details of all cultivated flax samples are included in supplementary table 1.

**Fig. 1.**
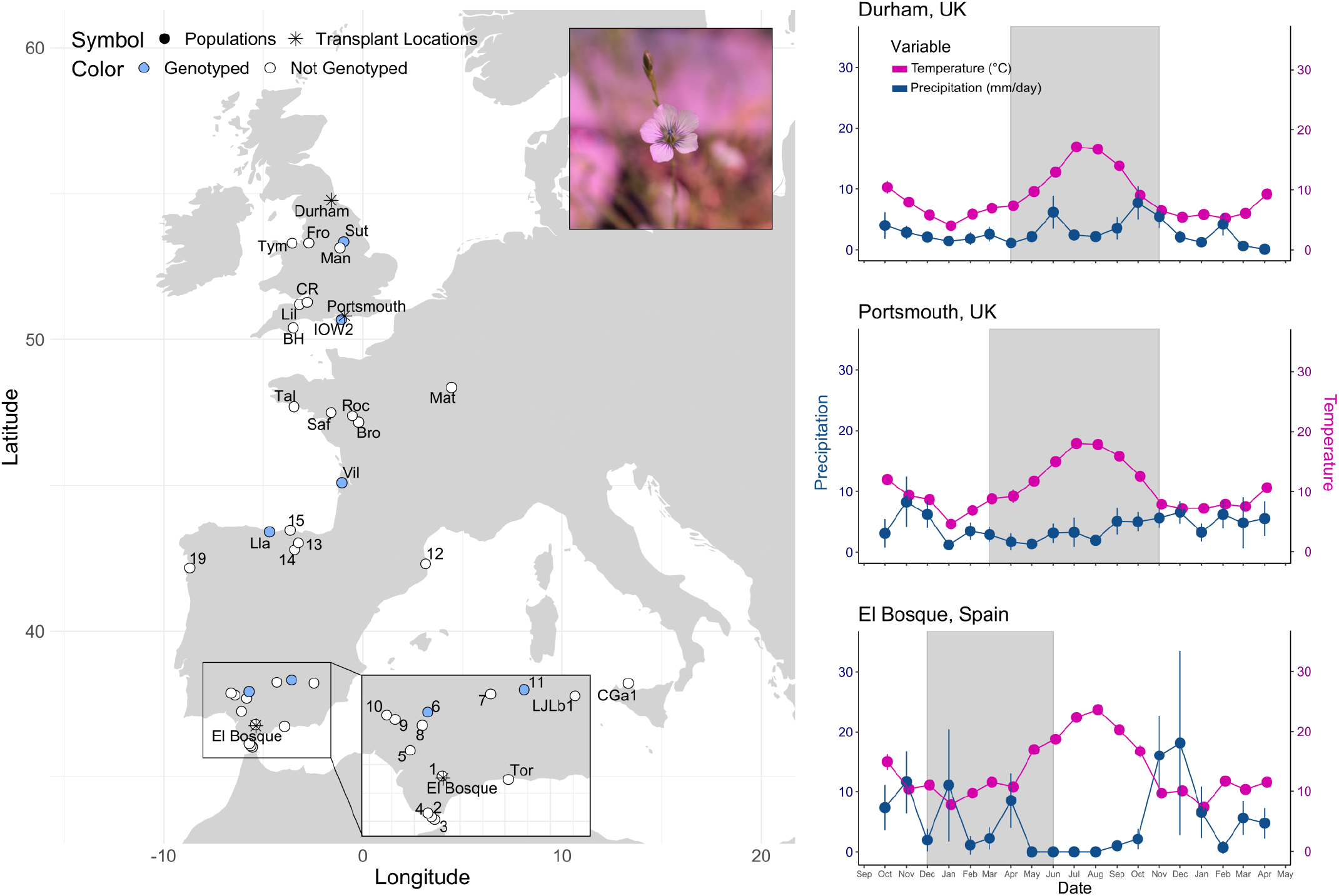
*Linum bienne* populations surveyed across its western distribution range (A), and the local climatic conditions that plants experienced during the reciprocal transplant experiment at the southern (El Bosque, Spain), central (Portsmouth, UK) and northern (Durham, UK) sites (1B), including temperature (pink) and precipitation (blue). In 1A (map), circles represent the populations surveyed, stars indicate reciprocal transplant sites (see Material & Methods Experiment 3), and blue circles indicate genotyped populations. In 1B (climate), the left and right y-axes are coloured in blue for precipitation and in pink for temperature.

### Description of historical climate of the populations surveyed

For all populations sampled in the field, data on mean monthly averages based on a 30-year period (1970 - 2000) for precipitation (mm), solar radiation (kJ/day*m2), average temperature (°C), minimum and maximum temperature (°C), vapour pressure (kPa), and wind speed (m/s) were retrieved using the WorldClim database at 30 arcsec resolution (Fick and Hijmans 2017). Climatic data were averaged by season. Months were assigned to seasons as follows: winter (December-February), spring (March-May), summer (June-August), and autumn (September-November). A principal component analysis (PCA) was used to reduce the dimensionality of the scaled climatic data and the first three principal coordinates (PCs) were retained since they explained most of the variation observed for each season. Pearson’s correlation coefficient was used to quantify the extent to which the first three principal components (hereafter climatic PC1, PC2 and PC3) were associated with latitude. All analyses included in this work were executed in R (R Core Team 2018) and all plots were produced with the R package ggplot2 v3.3.3 (Wickham 2016) together with Inkscape v1.1.1 (Inkscape Project 2020).

### Experiment 1: Population variation in flowering initiation under greenhouse conditions

The experiment was conducted in the insect free greenhouse facility of the University of Portsmouth. An average of 18 families from 32 populations were sown in October 2017 over a period of a week using seed collected from the wild (supplementary table 1 and 2). Before sowing, seeds were imbibed in gibberellic acid for 48 hrs to synchronise germination time. Sowing was randomized across populations and days so that the same set of populations was sown each day. Five seeds from the same family were placed in pots (9×9×10cm) with a mix soil and perlite 3:2 (the number of seedlings per pot did not affect the results of the analyses, results not shown). The positioning of populations and families was randomized across the greenhouse. Plants were grown using LED lights (14 hrs light at 22°C, 10 hrs dark at 16°C) throughout the experiment. Irrigation was done three times a week using an ebb and flow system and a commercial fertilizer was applied every two months by spraying plants. Since *L. bienne* is susceptible to powdery mildew (*Oidium* spp.), pest control was done using a sulphur burner. All plants were moved outside the greenhouse during summer months (from June to September) to avoid heat stress due to the absence of an automatic cooling system in the greenhouse facility and returned to the facility in September. The experiment was concluded in October 2018. Seedling emergence was monitored twice a week for two months. Adult plants were inspected three days a week to score flowering initiation, which was measured as the number of days between sowing date and first flowering for each family. During the period when plants remained outside the greenhouse facility, insect visitation was low (Perez-Barrales, personal observation) and, since autonomous self-pollination occurs upon flower opening, it was considered that outcross pollination was negligible. Cultivated flax has been estimated to be >95% selfing with gene flow <2% over 10 cm distance in the field (Jhala *et al*. 2011). Therefore, seed production was considered as derived from self-pollination and fruits were collected into separate paper envelopes per family as they ripened to represent the *F*_*1*_ generation used in Experiments 2 and 3.

Pearson’s correlation coefficient was used to investigate the association between flowering initiation against latitude, climatic PC1, PC2, and PC3. Linear regression models were used to describe the slope between flowering initiation (using population means) with latitude and climatic PC1 only, since correlation with PC2 and PC3 was negligible. Model selection was used to identify which of latitude or climatic PC1 better predicted flowering initiation by comparing all possible combinations of latitude and climatic PC1 (only one predictor, additive effect model, and additive effect model with the interaction term PC1 x Latitude) using ΔAICc with AICcmodavg R package v1.25 (Mazerolle 2012). A general linear model with binomial distribution was then used to describe the relationship between the proportion of families that flowered in each population with latitude.

### Experiment 2: Flowering initiation in response to vernalization

The *F*_*1*_ derived from 29 wild populations was used to test the effect of vernalization on flowering time (supplementary table 1 and 2). In total, 103 families (1 to 8 families per population) and 16 cultivars were exposed to vernalization and no vernalization conditions. Five seeds for each family and crop variety were sown in pots (10×9.5×10cm) filled with a mix of soil and perlite 3:1 and grown in controlled environment chambers (Models A3655 and A3658, Weiss-Gallenkamp, Loughborough, UK). Families (*L. bienne*) and cultivars (*L. usitatissimum*) were arranged in a randomized block design, replicating each treatment across two growth chambers each. The experimental conditions used were similar to those tested previously for *L. bienne* (Gutaker 2014). All seeds were cold-stratified for 3 days in darkness at 4°C, followed by 10 days at 22-20°C for 16:8 h in light:dark photoperiod to synchronize germination. Then, during 40 days, half the plants were exposed to vernalization conditions (4°C, 16:8 h light:dark photoperiod), while the rest were kept at 24-16°C and 16:8 h light:dark photoperiod (no vernalization conditions). When the vernalization treatment finished, plants from both treatments were kept in the same conditions (24-16°C, 16:8 h light: dark photoperiod). Seed germination was recorded for two weeks, survival was monitored until blooming, and plant size was quantified as the height and number of basal branches at the first open flower. Flowering initiation was measured as the number of days from sowing until the first flower opened in each pot with regular monitoring until the end of the experiment after 320 days. Linear mixed effect models were used to investigate flowering initiation responses to vernalization across populations for *L. bienne* and cultivars in *L. usitatissimum* (R package *lme4* v1.1-7, Bates *et al*. 2015). The model for *L. bienne* included the fixed factors; population, treatment and the interaction effect, while family and block were random factors. For *L. usitatissimum*, the model included treatment, cultivars and the interaction term as fixed factors, and block as random factor. The p-values were calculated based on Satterthwaite’s methods (lmertest, Kuznetsova *et al*. 2017; RLRsim, Scheipl *et al*. 2008). For both species, Pearson’s correlation coefficient was calculated to examine the association of flowering initiation with plant size in order to describe the level to which development interacts with flowering under the two vernalization treatments.

Vernalization sensitivity was used to compare the magnitude of the response to vernalization in wild and cultivated flax. Environmental sensitivity was calculated as the difference between the measurement of a family in different environments, representing a measure of the plastic response of the genotype (Falconer 1990). In the present study, vernalization sensitivity was calculated as:

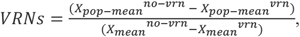

where *X*_*pop-mean*_^*no-vrn*^ and *X*_*pop-mean*_^*vrn*^ represent the population mean (or cultivar mean) values for the no-vernalization and vernalization treatment, while *X*_*mean*_^*no-vrn*^ and *X*_*mean*_^*vrn*^ represent the mean value of the no-vernalization and vernalization treatment, respectively. Sensitivity values range from positive to negative: positive values imply an earlier flowering initiation while negative values represent a delay in flowering initiation in vernalized plants respectively. Values closer to zero indicate a less sensitive response to vernalization cues. Sensitivity values between *L. bienne* and *L. usitatissimum* were compared with a t-test using bootstrapping (10.000 permutations) to calculate the significance of the difference using the R package MKinfer v0.6 (Kohl 2020) Finally, model selection was used to investigate if vernalization sensitivity of *L. bienne* populations was better predicted by latitude of origin or PC1 as described in Experiment 1, and linear models were also used to describe the slope of the relationship of vernalization sensitivity with latitude of origin and PC1.

### Experiment 3: Variation in population performance under reciprocal transplant field conditions

The field experiment was conducted at three sites representing the southernmost (El Bosque, Spain) and northernmost (Durham, UK) extremes of the western latitudinal range, and a central part of the latitudinal range of *L. bienne* (Portsmouth, UK; see Fig. 1A). The experiment was conducted using the *F*_*1*_ seed of 31 populations (2 to 10 families per population, supplementary table 3) which were replicated across sites, using a total of 300 pots per site, simulating provenance trials (see Anderson *et al*. 2015; Anderson and Wadgymar 2020). Pots were distributed in three blocks, with populations represented in two to three blocks per site and families distributed at random within blocks (supplementary table 3). Between 5 and 10 seeds were sown in each pot using peat-based compost. Since El Bosque represents the southernmost location, where droughts start at the end of spring until the start of the autumn (Fig. 1B), plants were sown in late November 2018 (23/11/2018). In Portsmouth (central location), plants were sown at the beginning of March 2019 (04/03/2019), while in Durham (northern location) sowing occurred three weeks later (26/03/2019). This strategy was chosen to account for differences in seasonality that the three locations experience (Fig. 1B) (for a similar approach see Toräng *et al*. 2015). After seedling emergence, a randomly selected plant per pot was tagged with a yellow pin to record flowering initiation, fruit production (number of fruits produced by plants) and plant survival weekly until the end of the growing season in October 2019. During the experiment, the local variation in temperature was monitored by recording hourly maximum and minimum temperature using a covered data logger at soil level. Daily precipitation and air temperature were retrieved using local weather stations (data retrieved from https://www.ogimet.com/synops.phtml.en, courtesy of R. Brugge). The weather conditions that the plants experienced during the experiment are summarised in Fig. 1B. Monthly precipitation was regular at the Portsmouth and Durham sites during the experiment, while plants at El Bosque experienced an intense drought from May, with a substantial increase of temperatures.

The analyses of data included general linear models of plant survival to assess whether probability of survival varied depending on distance from homesite across the transplant locations. The probability of survival was analysed separately for two different phenological moments, including ca. 100 days after the start of the experiment to represent plant survival right before the start of flowering (hereafter survival before flowering), and at the end of the growing season in autumn (hereafter survival to the autumn). The latter was analysed using the data retrieved from Portsmouth and Durham since no plants survived the summer drought at El Bosque. The analyses were done by pooling the data by population due to the variation in survival, flowering and fruiting among at family level (supplementary table 3). The survival analyses included a binomial distribution with logit link function, including distance from homesite, transplant location, and their interaction as predictors. The contrasts for transplant location were set to sums so that the model intercept could be interpreted as the grand mean for a trait over all transplant locations. Because of high mortality at El Bosque and limited flowering at Durham (see results below), the analyses on flowering initiation and fruit production were done using the data retrieved at Portsmouth. Variation in flowering initiation was analysed in relation to latitude and PC1 as described in Experiment 1 and 2. For fruit production, since the number of families with fruits varied by population, the average number of fruits produced by population was used as response variable in the GLM including Poisson distribution with log link and distance from homesite as predictor. Moreover, a similar GLM model was applied to fruit number using flowering initiation as predictor (instead of distance from homesite) to assess whether flowering initiation was under selection in Portsmouth. The R package emmeans v1.5.0 (Lenth, 2020) was used to assess post-hoc differences between sites for survival before flowering.

### Flowering initiation comparisons across all experiments

Pearson’s correlation coefficient was used to investigate the robustness of flowering initiation responses across different environments (Experiment 1 to 3) and generations to determine the fixed genetic influence of population of origin. For experiment 3, the flowering initiation data used corresponded to the data generated at the Portsmouth experimental site.

### Population genotyping and population genetic structure

To explore whether the phenological patterns observed might be due to underlying population structure, families of six populations from different latitudes were genotyped (Fig. 1A, supplementary table 1). Genotyping was conducted as described in Landoni et al. (Landoni *et al*. 2020) using microsatellite markers developed *ad hoc* for population genetic studies in *L. bienne*. Several of the 16 SSR loci developed showed signs of duplication, therefore each allele at each locus was considered as a dominant present or absent locus per se, giving a total of 64 dominant loci. Three of these dominant loci were excluded from further analyses because they were fixed across all individuals. The genetic structure of the six populations was investigated using Bayesian clustering with the software STRUCTURE v.2.3.4 and the StrAuto pipeline (Pritchard *et al*. 2000; Chhatre and Emerson 2017). For Bayesian clustering, we applied the admixture model following Hardy-Weinberg equilibrium and correlated allele frequencies. Run parameters included 10 independent replications with a burn-in period of 50,000 followed by 1,000,000 Markov chain Monte Carlo iterations with K number of genetic clusters varying from one to six. To determine the optimal number of clusters, we calculated the ad hoc measure *Δ*K, according to Evanno et al. (Evanno *et al*. 2005). In addition, discriminant analysis of principal components (DAPC) was performed with the R package *adegenet* v2.1.3 (Jombart *et al*. 2010). For the DAPC analysis, the optimal number of clusters and individual probabilities of assignment to clusters was identified using find.clusters() following dapc() using the R package adegenet v2.1.3. The genetic clustering results were plotted with the aid of R package pophelper v2.3.0 (Francis 2016).

## Results

### Description of historical climate of the populations surveyed

Following principal component analysis of climatic variables, the first principal component (PC1) explained 63.7% of the variance, with larger positive loadings representing vaporization, temperature radiation and vaporization, followed by winter precipitation, while the larger negative loadings corresponded to summer precipitation, followed by winter and autumn winds. The second principal component (PC2) explained 18.8% of the variance, with large positive loadings for wind and vaporization, winter minimum temperature, and precipitation during autumn and winter. Negative PC2 loadings represented maximum temperatures during spring and summer. The third principal component (PC3) explained 10% of the variance, with a larger positive loading corresponding to wind. The larger negative loading corresponded to precipitation from spring to winter. A summary of the results of the PCA analysis is included in supplementary table 4.

### Experiment 1: Population variation in flowering initiation under greenhouse conditions

Seedlings from 779 families out of 854 emerged within a month, with an average of three plants per pot. In total, 442 families flowered during the one-year experiment and plants that did not flower were still alive by the time the experiment was concluded. Plants started to flower three months after sowing (120 days), but on average flowering started 190 days after sowing (supplementary table 2). Correlations with flowering initiation were statistically significant for latitude (r = 0.70, p < 0.0001) and climatic PC1 (r = -0.78, p < 0.0001), but not for PC2 (r = 0.06, p > 0.05) and PC3 (r = -0.02, p > 0.05). Hence, PC2 and PC3 were excluded from further analyses. The linear regression model showed that flowering initiation was delayed ca. 3 days per latitude degree going north (Table 1). Using model selection based on ΔAICc, it was found that the model explaining most variation in flowering initiation (50% based on adjusted R-squared) only included the climatic PC1, as opposed to models with only latitude, or with both latitude and climatic PC1 (Table 1). The relationship between flowering initiation and PC1 was negative so that flowering initiation advanced around 5 days for every increased unit of climatic PC1 (Table 1), with climatic PC1 corresponding to increasing temperature, increasing radiation, and decreasing summer precipitation (supplementary table 4). The relationship between the proportion of families that flowered with latitude was negative (estimate ± s.e., intercept = 7.79 ± 0.65, slope = - 0.17 ± 0.01, Chi square = 162.1, d.f. = 1, P < 0.001) so that, at the population level, less families from northern latitudes flowered within the duration of the experiment.

**Table 1.** Model selection to compare flowering initiation determined by latitude and climate summary in Experiment 1 (greenhouse), Experiment 2 (vernalization) and Experiment 3 (reciprocal transplant experiment) in *Linum bienne* populations. The results of model selection are included for the models tested (univariate models with only latitude or PC1, additive model, a model with the interaction term, and the null model with only the intercept), including the number of terms of the model (K), AICc values, the difference with regards the model with the lowest value (Delta AICc) and the parsimony weighting of support comparing these models (Weights AICcWt). Also shown are the summary results including: the intercepts, slopes, p-values, residual degrees of freedom (df.res), *r*^*2*^, and adjusted *r*^*2*^, of the univariate linear models between days to flowering initiation (Experiment 1 and 3) and vernalization sensitivity (Experiment 2) with latitude and PC1. The selected models are in bold.

| Experiment | Response Variable | Model Selection |  |  |  |  | Model | Predictor | Estimate | p-value | df.res | R2 | Adj-R2 |
| --- | --- | --- | --- | --- | --- | --- | --- | --- | --- | --- | --- | --- | --- |
|  |  | Predictors | K | AICc | Delta_AICc | Weights AICcWt |  |  |  |  |  |  |  |
| 1 - Greenhouse | Flowering Initiation (days from sowing) | **PC1** | 3 | 280.2 | 0 | 0.61 | **PC1** | Intercept | 186.00 | <0.0001 | 30 | 0.61 | 0.60 |
|  |  | PC1*Latitude | 5 | 282.6 | 2.37 | 0.19 |  | PC1 | -5.04 | <0.0001 |  |  |  |
|  |  | PC1+Latitude | 4 | 282.6 | 2.4 | 0.18 |  |  |  |  |  |  |  |
|  |  | Latitude | 3 | 288 | 7.79 | 0.01 | Latitude | Intercept | 43.03 | n.s. | 30 | 0.50 | 0.48 |
|  |  | Null | 2 | 307.8 | 27.58 | 0 |  | Latitude | 3.37 | <0.0001 |  |  |  |
| 2 - Vernalization | Vernalization Sensitivity | **PC1** | 3 | 56.62 | 0 | 0.54 | **PC1** | Intercept | 1.06 | <0.001 | 23 | 0.27 | 0.24 |
|  |  | Latitude | 3 | 58.25 | 1.62 | 0.24 |  | PC1 | -0.09 | <0.01 |  |  |  |
|  |  | PC1 + Latitude | 4 | 59.41 | 2.79 | 0.13 |  |  |  |  |  |  |  |
|  |  | PC1 * Latitude | 5 | 61.39 | 4.76 | 0.05 | Latitude | Intercept | -1.64 | n.s. | 23 | 0.22 | 0.19 |
|  |  | Null | 2 | 61.9 | 5.28 | 0.04 |  | Latitude | 0.06 | <0.05 |  |  |  |
| 3 - Reciprocal Transplant | Flowering Initiation (days from sowing) | **PC1\*Latitude** | 5 | 233.9 | 0 | 0.36 | **PC1** | Intercept | 110.66 | <0.0001 | 28 | 0.34 | 0.31 |
|  |  | **PC1** | 3 | 234 | 0.1 | 0.34 |  | PC1 | -1.7281 | <0.0001 |  |  |  |
|  |  | Latitude | 3 | 235 | 1.03 | 0.21 |  |  |  |  |  |  |  |
|  |  | PC1+Latitude | 4 | 236.7 | 2.72 | 0.09 | Latitude | Intercept | 57.57 | <0.001 | 28 | 0.32 | 0.29 |
|  |  | Null | 2 | 243.9 | 9.93 | 0 |  | Latitude | 1.24 | <0.01 |  |  |  |

### Experiment 2: Flowering initiation in response to vernalization

At *L. bienne* family level, germination occurred in 84 out of 103 vernalized and in 89 out of 103 families in the non-vernalization treatment. A total of 93 families (254 plants) flowered, 82 (151) and 69 (103) for vernalization and no vernalization treatments respectively. However, by the end of this experiment, three families and 40 plants from 35 families had not flowered in the no vernalization treatment. The linear mixed effect model explained 82% of the variation in flowering initiation, showing statistically significant differences at the level of treatment, population, and the interaction term (Table 2). After vernalization, individual flowering initiation ranged between 84 to 143 days (mean±SD, n: 104.7±10.88, 151), whereas in the no vernalization treatment it ranged between 63 to 320 days (176.67 ± 64.71, 103; supplementary table 2). Hence, vernalization reduced days to flowering initiation by at least 72 days on average compared to the no vernalization treatment. The differences in population response to the vernalization treatment are represented by the reaction norm slopes (Fig. 2A), highlighting large differences between populations in the no-vernalization treatment. For example, the vernalization resulted in a mild delay in the start of flowering in some populations (i.e., TOR, IOW2, 5) with flowering occurring between 4 and 12 days later after vernalization, whereas other populations (i.e., IOW1, SAF, LLA) experienced a strong reduction in flowering initiation by more than 130 days after vernalization (supplementary table 2). In *L. usitatissimum*, germination occurred in all the cultivars used in the experiment in the vernalization and no vernalization treatment respectively. A total of 97 plants flowered, 47 in the vernalization treatment and 51 in the no vernalization treatment. About 73% of the variation in flowering initiation was explained by the linear mixed effect model, with significant effects by treatment, crop varieties, and the interaction effect (Table 2). Compared to *L. bienne*, flowering initiation in *L. usitatissimum* crop varieties occurred earlier in both treatments: vernalized plants flowered between 86 to 103.75 days (97.72 ± 7.15, 6), while non-vernalized plats flowered more rapidly between 56.25 to 110.5 days (70.31 ± 18.63.15, 51) (Fig. 2B, supplementary table 3). Pearson’s coefficient of correlation between flowering initiation and plant size are presented in Table 3. For *L. bienne*, the correlation with plant height was significant in the two treatments but much larger for the plants exposed to no vernalization conditions. In contrast, the correlation for number of basal stems was only significant for the no vernalization treatment. For *L. usitatissimum* plant size did not show a significant correlation with flowering initiation in either treatment.

**Table 2.** Linear mixed models to test the effect of vernalization on flowering initiation (measured as days from bolting) in *Linum bienne* populations and *Linum usitatissimum* cultivars. The summary results include the estimates for the intercept and predictor, confidence intervals, standard errors (S.E.), t-value and p-values.

| *L. bienne* |  |  |  |  |  | *L. usitatissimum* |  |  |  |  |  |
| --- | --- | --- | --- | --- | --- | --- | --- | --- | --- | --- | --- |
| Predictors | Estimates | Confidence Interval | S.E. | t-value | p-value | Predictors | Estimates | Confidence Interval | S.E. | t-value | p-value |
| Intercept | -102.4 | -132.24, -72.57 | 15.22 | -6.73 | <0.001 | Intercept | -35.15 | -62.57, -7.73 | 13.7 | -2.57 | 0.012 |
| Population | 2.1 | 1.88, 2.31 | 0.11 | 18.76 | <0.001 | Cultivar | 1.28 | 0.96, 1.61 | 0.16 | 7.79 | <0.001 |
| Treatment | 185.88 | 150.78, 220.98 | 17.91 | 10.38 | <0.001 | Treatment | 100.96 | 59.72, 142.24 | 20.65 | 4.89 | <0.001 |
| Population × Treatment | -1.94 | -2.20, -1.68 | 0.13 | -14.77 | <0.001 | Cultivar × Treatment | -0.88 | -1.38, -0.39 | 0.25 | -3.59 | 0.001 |
| **Random Effects** |  |  |  |  |  | **Random Effects** |  |  |  |  |  |
| σ<sup>2</sup> | 550.84 |  |  |  |  | σ<sup>2</sup> | 111.01 |  |  |  |  |
| τ<sub>00</sub> family | 133.66 |  |  |  |  | τ<sub>00</sub> block | 14.27 |  |  |  |  |
| τ<sub>00</sub> block | 4.12 |  |  |  |  | ICC | 0.11 |  |  |  |  |
| ICC | 0.2 |  |  |  |  | N block | 18 |  |  |  |  |
| N family | 96 |  |  |  |  | Observations | 97 |  |  |  |  |
| N block | 20 |  |  |  |  | Marginal-R<sup>2</sup>/<br>R<sup>2</sup> conditional | 0.69/ 0.73 |  |  |  |  |
| Observations | 253 |  |  |  |  |  |  |  |  |  |  |
| Marginal-R<sup>2</sup>/<br>R<sup>2</sup> conditional | 0.78/0.82 |  |  |  |  |  |  |  |  |  |  |

**Table 3.**
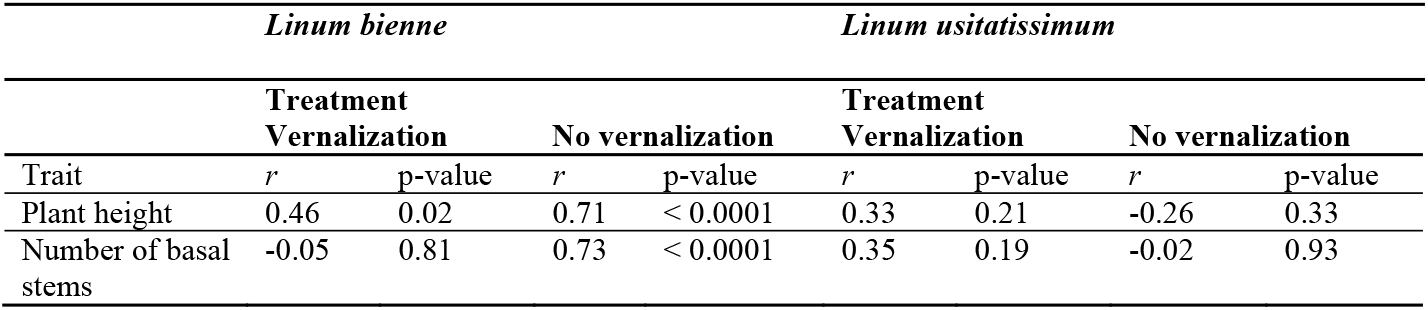
Pearson’s coefficient of correlation (r) and associated p-values of days to flowering initiation with plant height and the number of basal stems of *Linum bienne* and *Linum usitatissimum* plants exposed to vernalization and no vernalization conditions in Experiment 2. The coefficients were calculated using the population mean value and cultivar mean value of the traits (N = 25 for *L. bienne*, N = 16 for *L. usitatissimum*).

**Fig. 2.**
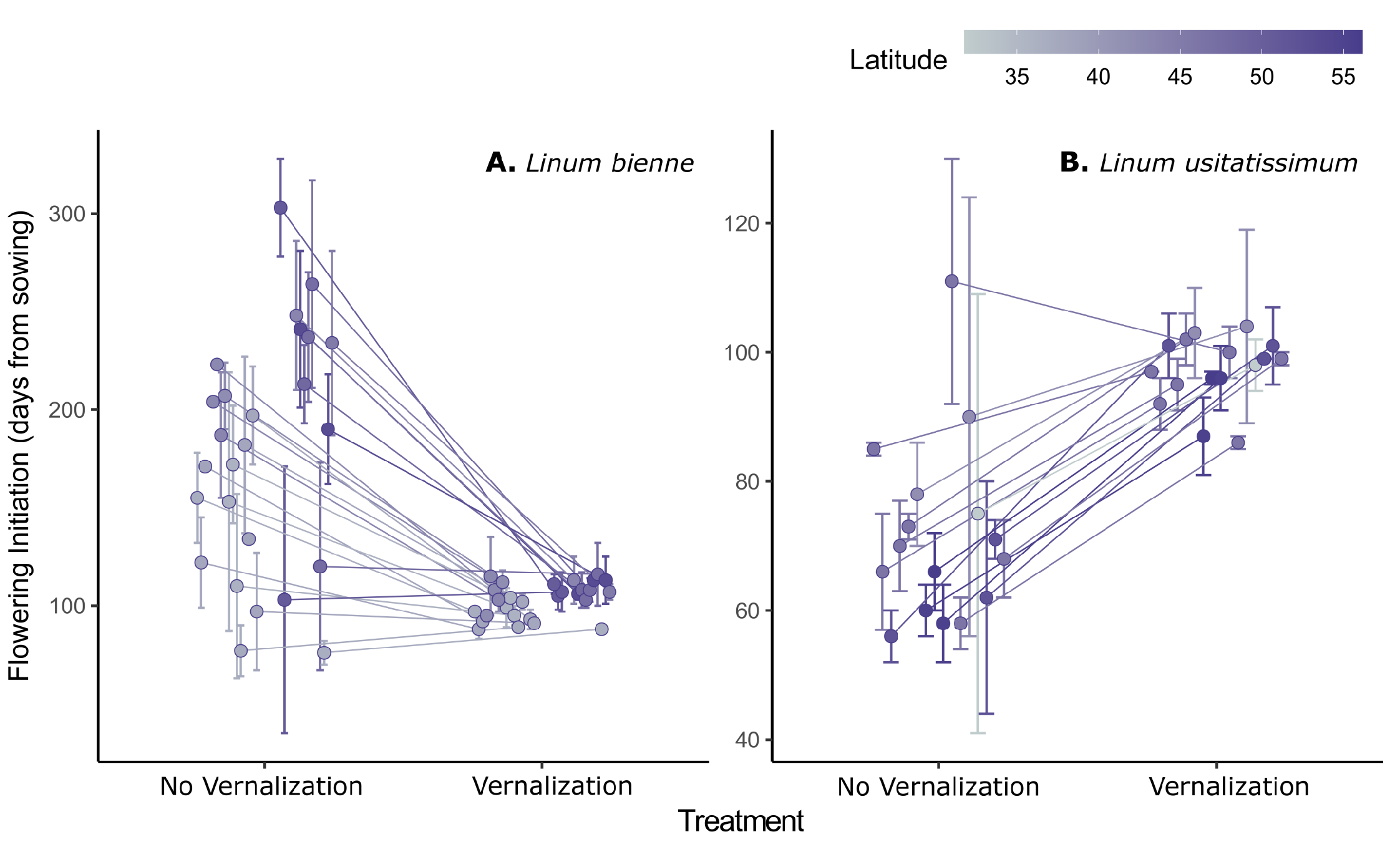
Reaction norm of days to flowering initiation in *Linum bienne* populations (A) and *Linum usitatissimum* varieties (B) in response to the vernalization manipulation. Points and error bars indicate mean and standard deviation of days to flowering initiation per population and treatment, and lines link populations across treatments. The colour scale represents the latitude of origin for *L. bienne* populations; for *L. usitatissimum*, the latitude was derived from the centroid coordinate of the country of origin for each cultivar

The magnitude of the vernalization response was reflected in sensitivity values, with statistically significant differences between the two species (t = 8.64, d.f.= 28.81, P<0.001). While values in *L. usitatissimum* were in general small and negative (−0.66± 0.33, 16, range: - 1.15, 0.29), vernalization sensitivity varied substantially among populations of *L. bienne* (1.96± 1.49, 26, range: -0.47, 5.15, supplementary table 2) from large positive values indicating a considerable reduction in days to flowering to a few populations exhibiting small negative values. The analysis of vernalization sensitivity in *L. bienne* showed negative relationship with latitude (Table 1). Following model selection and ΔAICc, it was found that the model that better predicted vernalization sensitivity only included PC1 (Table 1). The negative relationship between vernalization sensitivity and PC1 suggests that populations displaying greater responses to vernalization (i.e., larger reduction in flowering initiation time) were associated with an overall colder climate and wetter summer (negative PC1 loadings), while populations less responsive to vernalization (i.e., with no advance or small delay in flowering in response to cold cues) were related to warmer climate (positive PC1 loadings).

### Experiment 3: Variation in population performance under reciprocal transplant field conditions

A total of 154 families were used per experimental site of which 124, 123 and 120 germinated and 118, 119 and 109 survived to just before flowering at El Bosque, Durham and Portsmouth sites respectively. By the end of the experiment, 112 families were still alive at Portsmouth, 86 at Durham and no plants survived at El Bosque (supplementary table 3). From May, plants at El Bosque experienced an intense drought (Fig. 1C) which caused high mortality, so that only 14 families from 6 populations flowered. At Durham, 10 families from 9 populations flowered, while 110 families that represented well the latitudinal range flowered at Portsmouth (supplementary table 3). The average probability of plant survival before flowering was similar across the three experimental sites (LR Chi-square = 4.13, d.f. = 2, P= 0.13), and differences appeared associated with population distance from home site (LR Chi-square = 4.58, d.f. = 1, P= 0.03) and the interaction term (LR Chi-square = 77.00, d.f. = 2, P < 0.001). At El Bosque, there was a positive relationship between probability of plant survival and population distance from home site, while the relationship was negative at the central and northernmost experimental sites (Fig. 3A, Table 4). The analyses of plant survival to the autumn rendered similar results (Transplant site: LR Chi-square = 1.31, d.f. = 1, P= 0.25; Population distance from home site: LR Chi-square = 56.39, d.f. = 1, P<0.001; interaction term: LR Chi-square = 4.46, d.f. = 1, P < 0.001), with plant survival displaying a negative slope at both experimental sites (Fig. 3B, Table 4). In both models, the slope at the Durham site was steeper than the slope at the Portsmouth site (both P < 0.05) showing that selection against distant populations was stronger at the northernmost site than the central site (Table 4). With regards the analyses of the flowering and fruit data generated at Portsmouth, model selection showed that both a model with the interaction term (Latitude*PC1) and a model with only PC1 better predicted flowering initiation in Portsmouth (Table 1). However, the model with only PC1 was preferred since it represented a simpler model with less terms that agreed with the results of Experiment 1 and 2. The linear models showed that the relationship of flowering initiation was positive with latitude and negative with climatic PC1 (Table 1). The relationship between fruit production and the distance from homesite was significant (LR Chi square = 13.37, d.f.= 1, P < 0.001, Fig. 3C), displaying a negative slope. Fruit production also displayed a negative and significant relationship with flowering initiation so that populations that started flowering earlier produced more fruits (LR Chi square = 11.66, d.f.= 1, P < 0.001, Fig. 3D). Hence, while populations that started to flower earlier probably had a longer season to flower and set fruits, fruit production was higher in populations geographically closer to the experimental site.

**Table 4.**
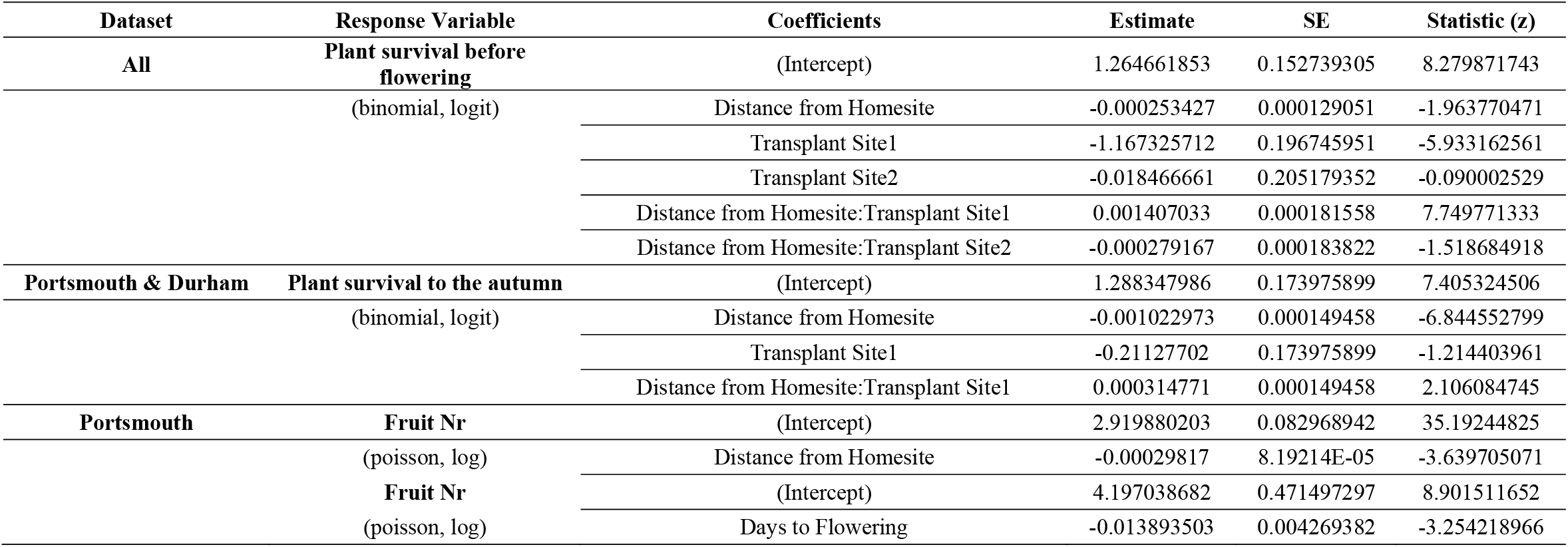
Estimates for the general linear models (glm) conducted on fitness components, including plant survival before flowering, plant survival to the autumn and fruit number (fruit nr), in response to distance from homesite. For fruit number, a glm was also conducted with days to flowering as predictor. The column “dataset” refers to the dataset used for the analyses included one (only Portsmouth), two (Portsmouth and Durham), or all three transplant locations (El Bosque, Durham, and Portsmouth). All models were conducted by setting contrasts to sum which affects how estimates are summed to obtain means for different levels of a factor.

**Fig. 3.**
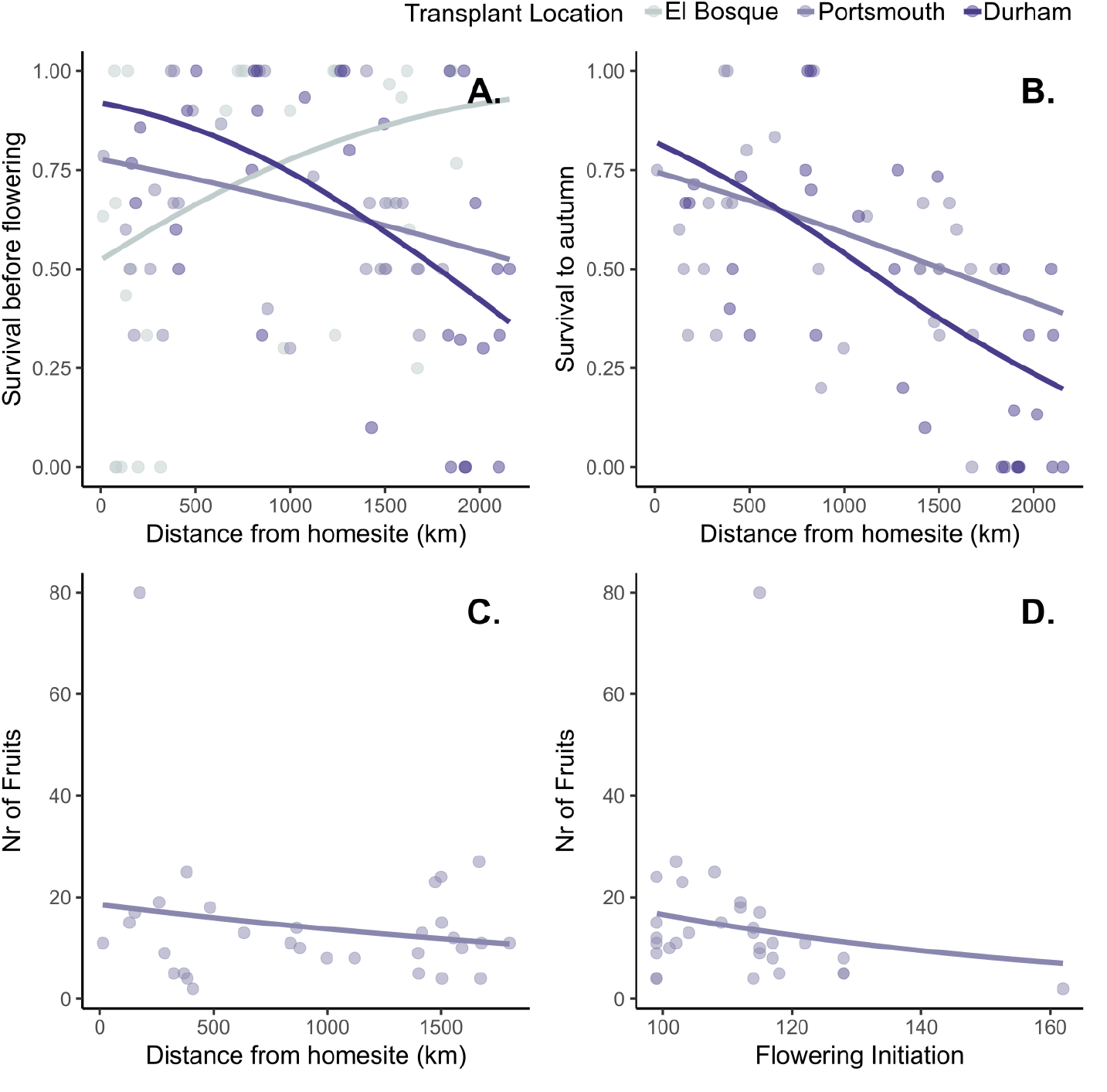
Performance of populations in the reciprocal transplant experiment along the western range of the distribution of *Linum bienne*. (A) Predicted probability of plant survival before flowering and (B) plant survival after the growing season (B) of *Linum bienne* populations exposed to the environmental conditions of El Bosque (southern site), Portsmouth (central site) and Durham (northern site), representing the conditions that the species experiences at the margin and at the centre of the distribution. None of the plants survived the summer drought at El Bosque so that survival after the growing season includes data from the sites at Portsmouth and Durham. (C) Relationship between mean fruit production per population and population distance from home site and (D) between mean fruit production per population and days to flowering initiation obtained at Portsmouth (central site) (the results ploted in C and D represent data only gathered at the Portsmouth site, see Results). In all figures, points represent population mean values coloured according to the transplant location, and lines represent the best fitting model predictions.

### Flowering initiation comparisons across all experiments

The correlation between flowering initiation generated with the F_0_ and F_1_ seeds (supplementary figure 1) was statistically significant and ranged between *r* = 0.51 (Experiments 1 and 3), *r* = 0.6 (Experiment 1 and Experiment 2 no vernalization treatment) and *r* = 0.7 (Experiments 1 and Experiment 2 vernalization treatment). The correlation of the F_1_ generation flowering initiation data was also statistically significant (supplementary figure 1), with values of *r* = 0.73 (Experiment 2 vernalization treatment and Experiment 3) and *r* = 0.53 (Experiment 2 no vernalization treatment and Experiment 3). This shows that the change in flowering initiation between F_0_ and F_1_ was similar across experiments, regardless the environment that F_0_ and F_1_ seeds experienced.

### Population genetic structure

Both Bayesian clustering and k-means clustering following DAPC identified the optimal number of genetic clusters to be two (supplementary figure 2A and 2B). The first cluster (southern cluster) contained populations 6 and 11 from southern Spain, while the second cluster (northern cluster) contained population LLA from northern Spain, population VIL from France, and populations IOW2 and SUT from UK (K = 2 in Fig. 4A). The Bayesian clustering assigned two individuals in population LLA to the southern cluster and identified a few admixed individuals between the southern and northern clusters for K = 2 or higher. This is potentially an indicator of gene flow from the southern to the northern cluster. For K greater than 2, the northern cluster was divided into the different populations it contained, with population LLA being the most distinct (Fig. 4A and 4B). Only three individuals from the population SUT were consistently assigned to the southern cluster across methods which might be an indicator of gene flow or sample mishandling.

**Fig. 4.**
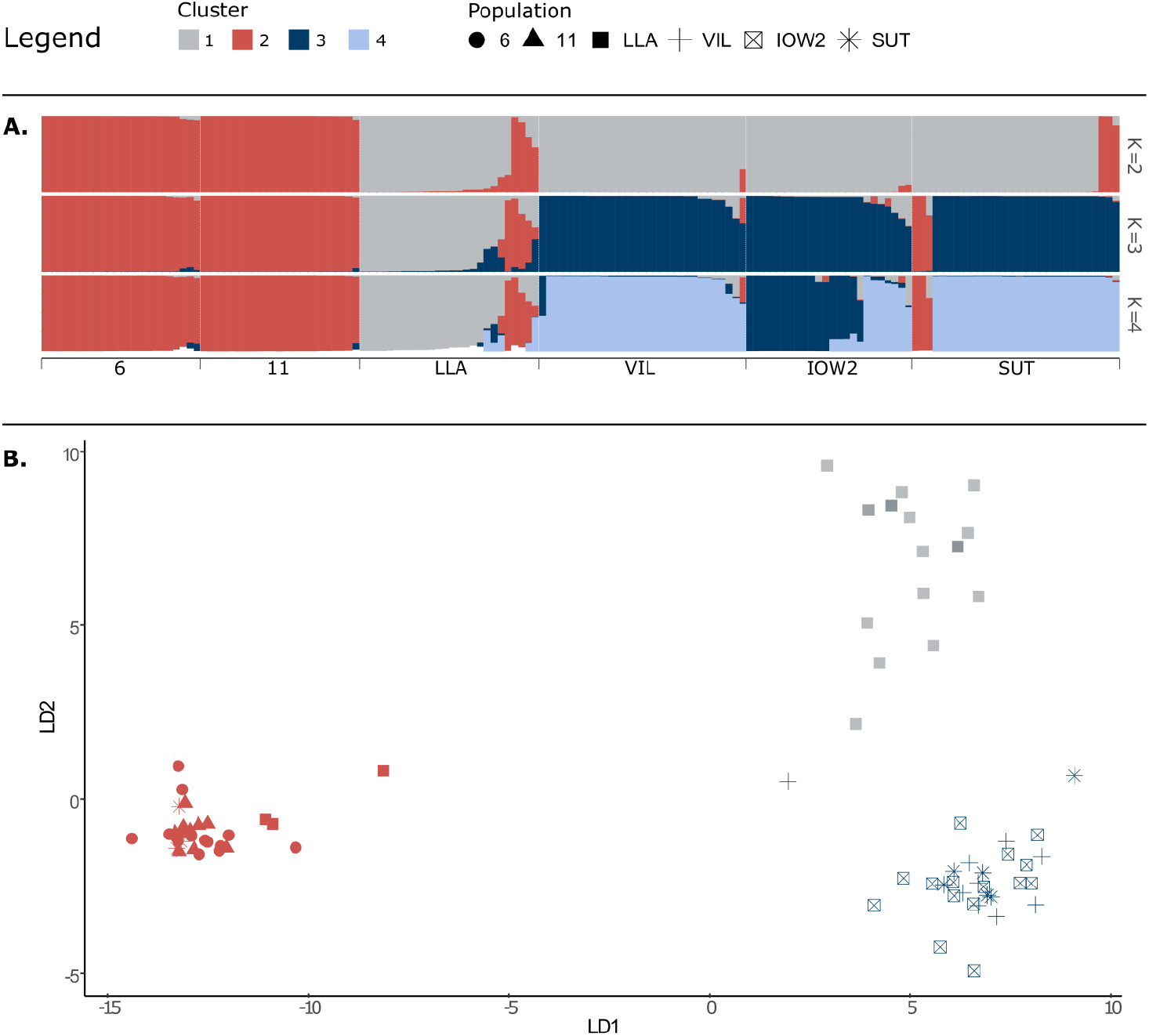
Genetic clustering of six *Linum bienne* populations where colours indicate clusters obtained with STUCTURE (A) or DAPC (B). (A) STRUCTURE results for two to four genetic clusters indicated with different colours are shown in each row starting from the top. Sample individuals are ordered along the x axis according to population. The colours of each vertical bar indicate the mean probabilities of assignment to each genetic cluster. (B) Discriminant analysis of principal components (DAPC) results showing samples separated according to the primary and secondary discriminant axes. Sample points are coloured according to the optimum number of genetic clusters (K = 3) and symbols indicate the populations.

## Discussion

### Population variation in flowering initiation

Latitudinal clines in flowering initiation are often considered the result of adaptation to climatic and environmental gradients to maximise plant growth and fitness in relation to the length of the growing season (Endler 1977; Boudry *et al*. 2002; Olsson and Ågren 2002; Stinchcombe *et al*. 2004; Keller *et al*. 2009; Colautti and Barrett 2013; Preite *et al*. 2015; Richardson *et al*. 2016). In its Western geographic range, *L. bienne* populations from the southern range started to flower earlier than populations from the northern range, with a decline in the probability of flowering in populations from northern latitudes. Flowering initiation was described using seeds collected in the field through different years. The consistent correlation among the data sets generated across experiments and between F_0_ and F_1_ suggests that maternal effects are minor in comparison with genetic effects. Since the data was generated under the controlled conditions of the greenhouse, it is probable that the latitudinal cline in days to flowering initiation emerged from a process of adaptation to environmental gradients, most likely climatic gradients, as evidenced by the association between flowering initiation and the climatic variables represented by PC1. Climate imposes strong selection on phenological traits, in turn creating genetic differentiation of populations and/or adaptation to local climatic conditions (Mendez-Vigo *et al*. 2011; Ågren and Schemske 2012; Burgarella *et al*. 2016; Lowry *et al*. 2019). The latitudinal gradient in flowering initiation described in this study agrees with the altitudinal differentiation observed for different phenotypic traits including flowering initiation among *L. bienne* accessions from Turkey (Uysal *et al*. 2012). Hence, the flowering initiation data in *L. bienne* provides additional evidence in support to the hypothesis that variation in phenology can be driven by environmental variation (Morente-López *et al*.; Endler 1977; Boudry *et al*. 2002; Stinchcombe *et al*. 2004; Franks *et al*. 2007; Keller *et al*. 2009; Mendez-Vigo *et al*. 2011; Colautti and Barrett 2013; Preite *et al*. 2015; Richardson *et al*. 2016; Lowry *et al*. 2019; Lampei *et al*. 2019; Thibault *et al*. 2020).

### Flowering initiation in response to vernalization

Vernalization is important to advance flowering in *L. bienne*, and the patterns described agree with those detected in species with similar geographic distributions (Exposito-alonso; Boudry *et al*. 2002; Burgarella *et al*. 2016). Most of *L. bienne* populations used in the experiment flowered without vernalization, but the manipulation reduced substantially the number of days to flowering after vernalization. Vernalization can also influence plant size and architecture (Stinchcombe *et al*. 2005; Mendez-Vigo *et al*. 2011; Adhikari *et al*. 2012; Quilot-Turion *et al*. 2013), some of the most important biomass yield parameters (Andrés and Coupland 2012; Lowry *et al*. 2019). Although some families flowered later or did not flower in absence of vernalization, the correlation between flowering initiation and plant size suggests that, in the absence of cold signals, plant development is important for flowering. Cold cues are used by plants to time flowering initiation late in winter or early in spring, providing an opportunity for seed maturation and dispersal in summer and germination in autumn. It is noteworthy that some of the *L. bienne* populations from the southern range did not advance flowering or slightly delayed flowering after vernalization. The life cycle of those populations could correspond to a different strategy. The significant population x treatment effect indicates that populations displayed considerable variation in the magnitude of the vernalization response. The population variation in response to the vernalization treatment probably reflects genetic differentiation for this trait and highlights local adaptive responses of flowering initiation to temperature cues (Boudry *et al*. 2002; Stinchcombe *et al*. 2005; Mendez-Vigo *et al*. 2011; Lewandowska-Sabat *et al*. 2012; Quilot-Turion *et al*. 2013). The responsiveness of flowering initiation has crucial implications for reproductive success in coping with local environmental heterogeneity (van Dijk and Hautekèete 2014; Blackman 2017; Prevéy *et al*. 2017).

The magnitude of the vernalization response was quantified as vernalization sensitivity to investigate environmental correlates. Consistent with Experiment 1, vernalization sensitivity followed a latitudinal cline that was better explained by a climatic gradient, with a negative relationship between vernalization sensitivity and climatic PC1 (Table1 & Fig. 2C). A small response or lack of response to cold cues is often interpreted as an adaptive strategy to escape hot and dry conditions during reproduction, typical of lower latitudes, or disturbances. By contrast, higher vernalization sensitivity might reflect adaptation to cold temperatures and shorter growing seasons of higher latitudes (Boudry *et al*. 2002). In the present study, populations representing the extremes of climatic PC1 exemplify both phenological responses from populations with low vernalization sensitivity (e.g., 4 in Spain) to deal with rainless summers typical of the Mediterranean climate of Southern Spain, to populations with large responses (e.g., MAN in UK) that experience longer winters and year-long precipitation in the UK (Fig. 1). Differences in vernalization sensitivity along climatic PC1 could contribute to the described intraspecific variability for life-histories in *L. bienne*. Testing this would require multi-year studies to observe survival and flowering initiation across multiple growing seasons. It would be of interest to determine the potential selective pressures driving differentiation in vernalization requirement as well as identify the genetic mechanisms underlying the vernalization responses (Boudry *et al*. 2002; Stinchcombe *et al*. 2005; Casao *et al*. 2011; Adhikari *et al*. 2012; Quilot-Turion *et al*. 2013; van Dijk and Hautekèete 2014; Höft *et al*. 2018; Thibault *et al*. 2020; Friedman 2020).

The incorporation of cultivated *L. usitatissimum* originating from across western Europe and Canada allowed us to establish comparisons with its wild progenitor and better understand the range of vernalization responses for flowering. Overall, *L. usitatissimum* flowered earlier that *L. bienne*, but the response to vernalization was opposite, with all varieties except for one delaying flowering, similarly to the response observed in some of the southern Spain populations with Mediterranean climate. The crop varieties displayed smaller sensitivity values and range of variation compared to its progenitor. Darapuneni et al. (Darapuneni *et al*. 2014) found that vernalization delayed flowering in winter varieties cultivated in Texas, but spring varieties of the Upper US Midwest and Canada were unaffected. Like other crops, vernalization in cultivated flax has probably been under selection to optimise flowering to different latitudes (Saisho *et al*.; Abbo *et al*. 2002). After its domestication in the Middle East, cultivated flax was progressively adopted in Europe which required adaptation to environments with strong seasonality in daylength and cold winters. This adaptation was mediated by secondary introgression of *L. bienne* into the gene-pool of cultivated flax, possibly introducing new variation at flowering time genes, favourable in northern latitudes (Gutaker *et al*. 2019). Further research on the genetic controls of vernalization to understand the adaptive nature of flowering initiation in *L. bienne* with latitude will be useful for breeding programs in a crop grown in a wide range of environments and climates (Sertse *et al*. 2019).

### Variation in population performance under reciprocal transplant field conditions

Reciprocal transplant experiments and provenance trials have demonstrated the adaptive significance of trait variation along environmental gradients to detect the magnitude of population maladaptation when exposed to new environments (Ågren and Schemske 2012; Colautti and Barrett 2013; Anderson *et al*. 2015; Toräng *et al*. 2015; Lowry *et al*. 2019; Anderson and Wadgymar 2020). In our experiment, populations exposed to the environmental conditions of the central (Portsmouth) and northern (Durham) sites showed a decline in the probability of plant survival (both before flowering and at the end of the growing season) with regards to the distance from home site. This agrees with the notion of population maladaptation under substantially new environmental conditions (Colautti and Barrett 2013; Anderson *et al*. 2015; Anderson and Wadgymar 2020). However, there was an initially positive relationship between survival before flowering and distance from homesite at the southernmost site (El Bosque), which indicates a lower performance of local populations at their home environment. Survival of local populations, which in this experiment represent margin populations at the southern range of *L. bienne*, could be compromised under contemporary environmental conditions (Sheth and Angert 2018; Midolo and Wellstein 2020; Anderson and Wadgymar 2020), with intensified droughts in the case of the Mediterranean region. Alternatively, an autumn planting date such as the one used in El Bosque might not be appropriate, and instead favour populations that are used to overwintering or to colder climates (i.e., northern populations). However, the few plants that survived the intense spring drought to flower were mostly from Spain, so the results at the southern site need to be interpreted with caution. Interestingly, most populations flowered at the central site, while only a few plants flowered at the southern and northern sites. This suggests that, together with vernalization, photoperiod might play an important role to initiate flowering in *L. bienne* (Samis *et al*. 2008; Lewandowska-Sabat *et al*. 2017), as found for some *L. usitatissimum* cultivars (Darapuneni *et al*. 2014). At the central site, days to flowering initiation showed a positive relationship with the latitude of origin, similarly to the results of Experiment 1 and 2, with an important effect of climate PC1 to predict flowering. The fact that flowering initiation displayed a negative relationship with fruit production, and that fruit production declined with distance from homesite supports the hypothesis of the adaptive nature of flowering initiation in *L. bienne*. The consistency of the results from Experiment 1 to 3, and the significant correlations of the flowering initiation data between F_0_ and F_1_ generations across the experiments highlight the strong genetic control of flowering initiation, which most likely emerged through local adaptation. The results of Experiment 3 are only partial due to high mortality and low flowering at the southern and northern sites respectively. Future field experiments should be optimised methodologically in order to understand the influence of environmental factors in the performance of populations to better estimate adaptation along the latitudinal cline.

### Genetic population differentiation

Our population genetic analysis only covered a subset of the populations included in the flowering initiation experiments, but genetic differentiation patterns suggest a strong divide in two groups represented by populations from southern Spain and by populations from northern Spain up to Northern England (Fig. 4A). For greater numbers of genetic clusters, it was possible to discriminate further populations suggestive of smaller scale genetic structure. Other plant species distributed across the Mediterranean basin and northern Europe show similar patterns (Hewitt 1999; Kadereit *et al*. 2005). This has been attributed to the shrinkage of their range to the Mediterranean area during the last glacial maximum (LGM) and a subsequent expansion to the north, from either the west or the east of the Mediterranean. Therefore, this result does not rule out the hypothesis that the clinal variation in flowering initiation in *L. bienne* is also determined by neutral processes (Keller *et al*. 2009). The strong genetic differentiation between populations from opposite margins of the distribution range is important for two reasons. First, southern populations are clearly distinct, both genetically (Fig. 4A and 4B) and phenotypically (early flowering) from northern populations so that patterns of local adaptation need to be also investigated within these genetic groups. Phenotypic and genetic differentiation attributed to local adaptation was found in other studies of *L. bienne*, but over smaller geographical scales (Uysal *et al*. 2010, 2012; Gutaker *et al*. 2019). Second, it raises the question whether re-colonization of northern Europe by *L. bienne* after the LGM occurred from Spain or from other refugia such as the South-East of the Mediterranean. *L. usitatissimum* was domesticated in the South-East and spread from there towards the North-West at first (Weiss and Zohary 2011; Fu 2012; Gutaker *et al*. 2019). Natural variation for agronomic traits available in south-western populations of *L. bienne* might be rare in the crop gene-pool if south-western populations of *L. bienne* remained relatively confined after the LGM and encountered the crop sometime after its domestication. Even for cultivated flax, the west of the Mediterranean is among the geographic regions that is less represented in worldwide collections, despite a recent study that found it harbours variation unique to the crop’s gene-pool (Sertse *et al*. 2019).

## Conclusions

This study revealed that flowering initiation and the response to vernalization in *L. bienne* co-vary with latitude in the west of the species’ native range. Flowering occurs later and is more responsive to vernalization for populations from increasing latitudes. The cline in flowering initiation and its vernalization response most likely reflects local adaptation to a range of environments from Mediterranean climate, with intense summer drought, to a Temperate Oceanic climate, with mild winters and wet summers. While local adaptation is a plausible driver of clinal variation in flowering, population structure and past demographic events should also be considered in future adaptation studies in *L. bienne* based on our findings that southern populations are genetically distinct from northern populations. Nonetheless, the latitudinal cline in flowering initiation remained consistent across different experiments indicating that flowering initiation is under strong genetic regulation. Additional experiments are required to confirm the adaptive nature of the vernalization response and its variation according to the temperature cue, photoperiod, or developmental stage of the plant. Further reciprocal-transplant studies, optimised to improve plant survival at the range extremes, should be conducted to investigate the extent that intraspecific variation and environmental factors determine lifespan and life-history in *L. bienne*, as variation in vernalization requirement (Experiment 2) and survival probability (Experiment 3) suggest. While the present study highlighted the importance of the vernalization pathway to reduce the days to flowering initiation, it is also of interest to investigate photoperiod and how it might interact with vernalization, in turn affecting flowering time and life-history across the wide geographic range. The variation described here also expands knowledge about flowering initiation in a crop wild relative, and the population collection we employed covers an unexplored section of the *L. bienne* native distribution along its whole latitudinal range. The patterns detected open new questions regarding what genetic mechanisms underpin the population differentiation of flowering in the species. This knowledge will be vital to predict plant phenological responses in a currently changing climate.

## Acknowledgements

The authors thank the support provided by Xavier Picó to conduct the reciprocal transplant experiment. B.L. was funded by a fellowship programme of the University of Portsmouth. P.S.M. was funded by a CONACyT (Mexico) Postdoctoral Research Fellowship and supported by Posgrado en Ciencias Biológicas of the Universidad Nacional Autónoma de México (UNAM). R.H.F.H. was funded by a UK BBSRC CASE PhD studentship BB/R506321/1. Fieldwork was conducted with fund provided by the Wild Flower Society to B.L. and by a travel grant from the Percy Sladen Memorial Fund to R.P.B.

## Data accessibility

Should the manuscript be accepted, the data supporting the results in the paper will be archived in Dryad and the data DOI will be included at the end of the article

