## supplementary for "Local climate and vernalization requirements explain the latitudinal patterns of flowering initiation in the crop wild relative *Linum bienne*"

SUPPLEMENTARY TABLES

Supplementary table 1. Surveyed populations of *Linum bienne* and cultivars of *Linum usitatissimum*, with indication of the type of species (wild for *L. bienne*, type of cultivar as oilseed or fibre for L. *usitatissimum*). The table includes number of families collected in the field, date of collection, country and locality, coordinates and elevation (metres above sea level), and cultivar provider. Column "Experiment" indicates what populations were used in Experiment 1, Experiment 2 and Experiment 3, and for genotyping (gt). For *L. bienne*, the seeds collected in the wild were directly used in experiments (experiment 1) or propagated in the greenhouse to generate an F1 (used in experiments 2 and 3). Populations were represented by different individuals (families). For *L. usitatissimum* all seeds represent a unique cultivar, and no distinction is done between individuals or families. Seeds provided by breeding companies and institutes were first propagated in the greenhouse for one generation before being used in Experiment 2. For *L. bienne*, coordinates represent the exact population's location while for *L. usitatissimum*, coordinates represent the centroid of the country of origin and no altitude is available.

| Species | Type | Population / Cultivar | Families Collected | Date of collection | Country | Locality / Provider | Lat | Lon | Alt | Experime nt |
|---|---|---|---|---|---|---|---|---|---|---|
| bienne | wild | 3 | 41 | 5/6/2016 | Spain | Guadalmesi | 36.036 | -5.556 | 186 | 1, 2, 3 |
| bienne | wild | 2 | 32 | 5/6/2016 | Spain | Virgen de la Luz Santuary Facinas | 36.081 | -5.626 | 41 | 1, 2, 3 |
| bienne | wild | 4 | 34 | 5/6/2016 | Spain | El Nene, Facinas | 36.151 | -5.705 | 36 | 1, 2, 3 |
| bienne | wild | Tor | 18 | 11/7/2013 | Spain | Torrox Costa, Malaga | 36.740 | -3.926 | 24 | 1, 2 |
| bienne | wild | 1 | 47 | 4/6/2016 | Spain | Llanos del Rabel trail | 36.800 | -5.393 | 624 | 1, 2, 3 |
| bienne | wild | 5 | 35 | 7/6/2016 | Spain | Puebla del Río-Aznalcazar, Sevilla | 37.259 | -6.097 | 12 | 1, 2, 3 |
| bienne | wild | 8 | 27 | 8/6/2016 | Spain | Road 432, Km 12 Road to El Pedroso | 37.702 | -5.832 | 128 | 1, 2, 3 |
| bienne | wild | 9 | 36 | 9/6/2016 | Spain | N-433 before exit to Zufre-La Granada de Riotinto, | 37.809 | -6.429 | 446 | 1, 2, 3 |

| | | | | | | | | | | |
|---|---|---|---|---|---|---|---|---|---|---|
| | | | | | | Huelva | | | | |
| bienne | wild | 10 | 28 | 9/6/2016 | Spain | Linares de la Sierra, Huelva | 37.882 | -6.618 | 519 | 1, 2, 3 |
| bienne | wild | 6 | 29 | 8/6/2016 | Spain | Constantina-Cazalla de la Sierra, Sevilla (trail) | 37.936 | -5.711 | 529 | 1, 2, gt |
| bienne | wild | CGa1 | 21 | 6/6/2017 | Italy | Capo Gallo, Sicily | 38.217 | 13.322 | 53 | 1, 3 |
| bienne | wild | LJLb1 | 15 | 15/6/2014 | Spain | Las Juntas, Jaen | 38.221 | -2.454 | 1325 | 1, 3 |
| bienne | wild | 7 | 21 | 8/6/2016 | Spain | Cardeña-Villa del Río, Cortijo Tejoneras, Córdoba | 38.253 | -4.317 | 752 | 1, 2, 3 |
| bienne | wild | 11 | 38 | 14/6/2016 | Spain | La Aliseda, Finca La Inmmediata (Km 3), Jaen | 38.331 | -3.581 | 710 | 1, 2, 3, gt |
| bienne | wild | 19 | 31 | 15/7/2016 | Spain | Universidad de Vigo | 42.171 | -8.684 | 483 | 1, 2 |
| bienne | wild | 12 | 51 | 17/6/2016 | Spain | Palau-Savereda | 42.310 | 3.152 | 105 | 1, 2, 3 |
| bienne | wild | 14 | 38 | 20/6/2016 | Spain | Tartales de Cilla | 42.794 | -3.425 | 670 | 1, 2 |
| bienne | wild | 13 | 35 | 19/6/2016 | Spain | Quincoces de Yuso-Relloso, Burgos | 43.029 | -3.240 | 741 | 1, 2, 3 |
| bienne | wild | Lla | 35 | 21/7/2014 | Spain | Llanes, Asturias | 43.407 | -4.688 | 26 | 1, 2, 3, gt |
| bienne | wild | 15 | 32 | 20/6/2016 | Spain | Cantabria-Carriazo.Galizano | 43.463 | -3.653 | 50 | 1, 2, 3 |
| bienne | wild | Vil | 38 | 2/7/2016 | France | Villeneuve, Charente Maritime | 45.094 | -1.050 | 21 | 1, 2, gt |
| bienne | wild | Bro | 39 | 4/7/2016 | France | Brossay, Maine et Loire | 47.167 | -0.207 | 69.5 | 1, 3 |
| bienne | wild | Roc | 40 | 4/7/2016 | France | Domaine de Rochambeau, Maine et Loire | 47.387 | -0.526 | 47.5 | 1, 2, 3 |
| bienne | wild | Saf | 30 | 5/7/2016 | France | Saffre, Loire Atlantique | 47.496 | -1.593 | 25 | 1, 2, 3 |
| bienne | wild | Tal | 40 | 6/7/2016 | France | Pointe du Talude, Morbihan | 47.700 | -3.455 | 14 | 1, 2, 3 |
| bienne | wild | Mat | 25 | 29/6/2016 | France | Mathaux, Aube | 48.357 | 4.459 | 125.5 | 1, 2 |
| bienne | wild | BH | 13 | 9/7/2017 | UK | Barry Head | 50.400 | -3.493 | 55 | 1, 3 |
| bienne | wild | Dor | NA | NA | UK | Dorset / Emorsgate Seeds | 50.600 | -2.010 | NA | 2 |
| bienne | wild | IOW2 | 42 | 24/9/2016 | UK | Bembridge, 2nd stop, Isle of Wight | 50.682 | -1.075 | 9 | 1, 2, gt |
| bienne | wild | IOW1 | 26 | 24/9/2016 | UK | Bembridge, 1st stop, Isle of Wight | 50.691 | -1.095 | 1 | 1, 2, 3 |
| bienne | wild | Lil | 10 | 9/7/2017 | UK | Lilstock | 51.202 | -3.187 | 23 | 1, 3 |
| bienne | wild | CR | 10 | 8/7/2017 | UK | Cheddar Reservoir | 51.278 | -2.795 | 15 | 1, 3 |
| bienne | wild | Man | 41 | 10/9/2016 | UK | Mansfield, Nottinghamshire | 53.137 | -1.144 | 124 | 1, 2, 3 |
| bienne | wild | Tym | 40 | 2/9/2016 | UK | Tyr Mawr Holiday Park, Denbighshire | 53.303 | -3.553 | 5 | 1, 2, 3 |

| | | | | | | | | | | |
|---|---|---|---|---|---|---|---|---|---|---|
| bienne | wild | Sut | 42 | 9/9/2016 | UK | Sutton Cum Lound, Nottinghamshire | 53.353 | -0.959 | 15 | 1, 2, 3, gt |
| usitatissimum subsp. mediterraneum | cultivated, oilseed | Raba 0189 | - | - | Morocco | IPK | 31.792 | -7.093 | - | 2 |
| usitatissimum subsp. caesium | cultivated, oilseed | Gisa | - | - | Italy | IPK | 41.872 | 12.567 | - | 2 |
| usitatissimum subsp. mediterraneum | cultivated, oilseed | Primus | - | - | Italy | IPK | 41.872 | 12.567 | - | 2 |
| usitatissimum subsp. elongatum | cultivated, fibre | Ariane | - | - | France | IPK | 46.228 | 2.214 | - | 2 |
| usitatissimum | cultivated, fibre spring | Aramis | - | - | France | Terre de Lin | 46.228 | 2.214 | - | 2 |
| usitatissimum | cultivated, fibre spring | Bolchoi | - | - | France | Terre de Lin | 46.228 | 2.214 | - | 2 |
| usitatissimum | cultivated, fibre spring | Eden | - | - | France | Terre de Lin | 46.228 | 2.214 | - | 2 |
| usitatissimum | cultivated, fibre winter | Olga | - | - | France | Terre de Lin | 46.228 | 2.214 | - | 2 |
| usitatissimum | cultivated, oilseed spring | Omegalin | - | - | France | Terre de Lin | 46.228 | 2.214 | - | 2 |
| usitatissimum | cultivated, oilseed winter | Volga | - | - | France | Terre de Lin | 46.228 | 2.214 | - | 2 |
| usitatissimum subsp. elongatum | cultivated, fibre | Blenda 04C | - | - | Netherlands | IPK | 52.133 | 5.291 | - | 2 |
| usitatissimum | cultivated, fibre | Suzanne | - | - | Netherlands | Flaxland | 52.133 | 5.291 | - | 2 |
| usitatissimum subsp. caesium | cultivated, oilseed | Tine Tammes Lila | - | - | Netherlands | IPK | 52.133 | 5.291 | - | 2 |
| usitatissimum subsp. elongatum | cultivated, fibre | Monarch | - | - | UK | IPK | 55.378 | -3.436 | - | 2 |
| usitatissimum subsp. caesium | cultivated, oilseed | Liral Crown | - | - | UK | IPK | 55.378 | -3.436 | - | 2 |
| usitatissimum | cultivated, oilseed | Marmalade | - | - | Canada | Flaxland | 56.130 | -106.347 | - | 2 |

Supplementary table 2. Summary results of the days to flowering initiation (mean, standard deviation and sample size) of the *Linum bienne* populations used to describe flowering initiation under the greenhouse conditions (Experiment 1, $F_0$ generation), and in the vernalization experiment (Experiment 2, $F_1$ generation) including the no vernalization (NV) and vernalization (V) treatment, as well as the vernalization sensitivity for *L. bienne* populations and *L. usitatissimum* cultivars. The summary includes the number of sown, emerged and flowered families used in both experiments.

| **Species** | **Population / Cultivar** | **Greenhouse Experiment (wild seed) Experiment 1** | | | | **Vernalization Experiment (F1 seed) Experiment 2** | | | | | **Vernalization sensitivity** |
|---|---|---|---|---|---|---|---|---|---|---|---|
| | | **Sown families** | **Emerged families** | **Flowered families** | **Flowering Start** | **Sown Families NV / V** | **Emerged Families NV / V** | **Flowered Families NV / V** | **Flowering Start No Vernalization** | **Flowering Start Vernalization** | |
| bienne | **1** | 30 | 20 | 20 | m = 149, sd = 21 | 6 / 6 | 4 / 2 | 4 / 2 | m = 155, sd = 23 | m = 97, sd = 2 (N = 4) | 0.8 |
| bienne | **2** | 30 | 29 | 29 | m = 137, sd = 23 | 2 / 2 | 2 / 2 | 2 / 2 | m = 153, sd = 66 | m = 99, sd = 10 | 0.75 |
| bienne | **3** | 29 | 20 | 12 | m = 189, sd = 29 | 4 / 4 | 4 / 4 | 4 / 4 | m = 172, sd = 30 | m = 104, sd = 3 | 0.94 |
| bienne | **4** | 27 | 22 | 20 | m = 139, sd = 21 | 5 / 7 | 5 / 5 | 5 / 5 | m = 110, sd = 47 | m = 95, sd = 5 | 0.21 |
| bienne | **5** | 30 | 24 | 23 | m = 139, sd = 18 | 2 / 2 | 2 / 2 | 2 / 2 | m = 77, sd = 13 | m = 89, sd = 2 | -0.18 |
| bienne | **6** | 29 | 24 | 20 | m = 190, sd = 22 | 3 / 3 | 3 / 3 | 3 / 2 | m = 182, sd = 45 | m = 102, sd = 4 | 1.11 |
| bienne | **7** | 21 | 19 | 15 | m = 197, sd = 23 | 1 / 1 | 1 / 0 | 1 / NA | m = 134, sd = NA | NA | NA |
| bienne | **8** | 24 | 22 | 18 | m = 176, sd = 31 | 4 / 4 | 2 / 2 | 2 / 2 | m = 197, sd = 25 | m = 93, sd = 5 | 1.45 |
| bienne | **9** | 30 | 27 | 25 | m = 129, sd = 18 | 3 / 3 | 3 / 2 | 3 / 2 | m = 97, sd = 30 | m = 91, sd = 1 | 0.09 |
| bienne | **10** | 28 | 28 | 26 | m = 175, sd = 29 | 5 / 5 | 4 / 4 | 4 / 4 | m = 122, sd = 23 | m = 88, sd = 5 | 0.47 |
| bienne | **11** | 30 | 30 | 16 | m = 179, sd = 36 | 1 / 1 | 1 / 1 | 1 / 1 | m = 171, sd = NA | m = 92, sd = NA | 1.1 |
| bienne | **12** | 30 | 29 | 10 | m = 209, sd = 24 | 1 / 1 | 1 / 1 | 0 / 1 | NA | m = 95, sd = NA | NA |
| bienne | **13** | 30 | 24 | 4 | m = 185, sd = 61 | 4 / 4 | 3 / 3 | 1 / 3 | m = 204, sd = NA | m = 115, sd = 20 | 1.23 |
| bienne | **14** | 8 | 3 | 1 | m = 195, sd = NA | 2 / 2 | 2 / 2 | 1 / 2 | m = 223, sd = NA | m = 108, sd = 5 | 1.6 |
| bienne | **15** | 31 | 31 | 12 | m = 200, sd = 24 | 7 / 5 | 5 / 5 | 4 / 5 | m = 187, sd = 32 | m = 103, sd = 6 | 1.17 |
| bienne | **19** | 30 | 29 | 20 | m = 217, sd = 23 | 6 / 6 | 3 / 1 | 2 / 1 | m = 207, sd = 17 | m = 112, sd = 6 | 1.32 |

| | | | | | | | | | | | |
|---|---|---|---|---|---|---|---|---|---|---|---|
| bienne | **BH** | 12 | 12 | 3 | m = 228, sd = 7 | - | - | - | - | - | - |
| bienne | **Bro** | 29 | 27 | 6 | m = 194, sd = 38 | - | - | - | - | - | - |
| bienne | **CGa1** | 21 | 21 | 21 | m = 137, sd = 21 | - | - | - | - | - | - |
| bienne | **CR** | 30 | 30 | 4 | m = 221, sd = 15 | - | - | - | - | - | - |
| bienne | **IOW1** | - | - | - | - | 3 / 3 | 3 / 3 | 2 / 3 | m = 303, sd = 25 | m = 105, sd = 7 | 2.75 |
| bienne | **IOW2** | 30 | 28 | 11 | m = 195, sd = 49 | 4 / 4 | 4 / 4 | 3 / 4 | m = 103, sd = 68 | m = 107, sd = 10 | -0.06 |
| bienne | **Lil** | 21 | 19 | 4 | m = 201, sd = 27 | - | - | - | - | - | - |
| bienne | **LJLb1** | 16 | 15 | 9 | m = 209, sd = 30 | - | - | - | - | - | - |
| bienne | **Lla** | 30 | 30 | 16 | m = 202, sd = 30 | 8 / 8 | 7 / 7 | 6 / 7 | m = 248, sd = 38 | m = 113, sd = 12 | 1.86 |
| bienne | **Man** | 30 | 28 | 8 | m = 213, sd = 39 | 5 / 5 | 4 / 4 | 3 / 4 | m = 241, sd = 40 | m = 106, sd = 3 | 1.87 |
| bienne | **Mat** | 25 | 25 | 22 | m = 208, sd = 21 | 5 / 5 | 5 / 5 | 5 / 5 | m = 213, sd = 20 | m = 108, sd = 9 | 1.46 |
| bienne | **Roc** | 29 | 29 | 16 | m = 213, sd = 29 | 1 / 1 | 1 / 1 | 1 / 1 | m = 237, sd = 33 | m = 103, sd = 4 | 1.86 |
| bienne | **Saf** | 27 | 26 | 7 | m = 186, sd = 37 | 4 / 4 | 4 / 4 | 3 / 4 | m = 264, sd = 53 | m = 108, sd = 6 | 2.17 |
| bienne | **Sut** | 29 | 29 | 8 | m = 208, sd = 24 | 1 / 1 | 1 / 1 | 0 / 1 | NA | m = 113, sd = 1 (N = 2) | NA |
| bienne | **Tal** | 28 | 26 | 9 | m = 208, sd = 30 | 4 / 4 | 4 / 4 | 1 / 4 | m = 120, sd = 53 | m = 116, sd = 16 | 0.05 |
| bienne | **Tor** | - | - | - | - | 1 / 1 | 1 / 1 | 1 / 1 | m = 76, sd = 6 | m = 88, sd = 2 | -0.16 |
| bienne | **Tym** | 30 | 28 | 8 | m = 224, sd = 10 | 4 / 4 | 4 / 4 | 2 / 4 | m = 190, sd = 28 | m = 113, sd = 12 | 1.07 |
| bienne | **Vil** | 30 | 30 | 21 | m = 205, sd = 31 | 4 / 4 | 4 / 4 | 3 / 4 | m = 234, sd = 47 | m = 107, sd = 4 | 1.77 |
| bienne | **Dor** | - | - | - | - | 3 / 3 | 2 / 3 | 0 / 2 | NA | m = 111, sd = 5 (N = 6) | NA |
| usitatissimum | **Aramis** | - | - | - | - | 2 / 2 | 2 / 2 | 2 / 2 | m = 85, sd = 1 | m = 97, sd = 0 | 0.44 |
| usitatissimum | **Ariane** | - | - | - | - | 4 / 4 | 4 / 4 | 3 / 4 | m = 66, sd = 9 | m = 92, sd = 4 | 0.96 |
| usitatissimum | **Blenda** | - | - | - | - | 4 / 4 | 4 / 4 | 4 / 4 | m = 56, sd = 4 | m = 101, sd = 5 | 1.62 |
| usitatissimum | **Bolchoi** | - | - | - | - | 2 / 2 | 2 / 2 | 2 / 2 | m = 70, sd = 7 | m = 95, sd = 4 | 0.89 |
| usitatissimum | **Eden** | - | - | - | - | 2 / 2 | 2 / 2 | 3 / 2 | m = 73, sd = 2 | m = 102, sd = 4 | 1.03 |
| usitatissimum | **Gisa** | - | - | - | - | 4 / 4 | 4 / 4 | 4 / 4 | m = 78, sd = 8 | m = 103, sd = 7 | 0.9 |
| usitatissimum | **Liral Crown** | - | - | - | - | 4 / 2 | 4 / 2 | 3 / 2 | m = 60, sd = 4 | m = 87, sd = 6 | 0.97 |
| usitatissimum | **Marmalade** | - | - | - | - | 2 / 2 | 2 / 2 | 2 / 2 | m = 66, sd = 6 | m = 96, sd = 1 | 1.08 |

| | | | | | | | | | | | |
|---|---|---|---|---|---|---|---|---|---|---|---|
| usitatissimum | **Monarch** | - | - | - | - | 4 / 5 | 4 / 5 | 4 / 4 | m = 58, sd = 6 | m = 96, sd = 5 | 1.37 |
| usitatissimum | **Olga** | - | - | - | - | 2 / 2 | 2 / 2 | 2 / 2 | m = 111, sd = 19 | m = 100, sd = 4 | -0.4 |
| usitatissimum | **Omegalin** | - | - | - | - | 4 / 2 | 4 / 2 | 4 / 2 | m = 58, sd = 4 | m = 86, sd = 1 | 1.04 |
| usitatissimum | **Primus** | - | - | - | - | 4 / 4 | 4 / 4 | 4 / 4 | m = 90, sd = 34 | m = 104, sd = 15 | 0.49 |
| usitatissimum | **Raba 0189** | - | - | - | - | 4 / 4 | 4 / 4 | 4 / 4 | m = 75, sd = 34 | m = 98, sd = 4 | 0.86 |
| usitatissimum | **Suzanne** | - | - | - | - | 4 / 2 | 4 / 2 | 4 / 2 | m = 62, sd = 18 | m = 99, sd = 0 | 1.36 |
| usitatissimum | **Tine Tammes Lila** | - | - | - | - | 4 / 4 | 4 / 4 | 4 / 4 | m = 71, sd = 3 | m = 101, sd = 6 | 1.11 |
| usitatissimum | **Volga** | - | - | - | - | 2 / 2 | 2 / 2 | 2 / 2 | m = 68, sd = 6 | m = 99, sd = 1 | 1.13 |

Supplementary table 3. Summary of the reciprocal transplant experiment conducted at El Bosque, Portsmouth and Durham, including the number of families and seeds per family used at each site, and the distance of the *Linum bienne* populations used in the experiment from home site in km. The table includes the number of pots, families, and seedlings that emerged, survived, and flowered during the experiment, as well as mean days to flowering per population, and mean number of fruits produced per population for Portsmouth only. NA indicates that those families that either did not emerge, flower or form fruits, depending on the column. For autumn survival (pots), NA indicates that for some pots within a population it was not possible to score survival (status unclear or pot moved by animals/weather), although for all populations it was possible to score whether at least one individual per family survived. Due to the large size (table not fitting the A4 dimension, the table is spitted in two parts.

**Part 1:**

| Transplant Site | Population | Latitude | Longitude | Distance from Homesite (km) | Sowing Date | Emergence Date | Total Pots | Total Families | Total Seeds | Emerged Pots | Emerged Families | Emerged Seedlings |
|---|---|---|---|---|---|---|---|---|---|---|---|---|
| El Bosque | **3** | 36.036 | -5.556 | 81.040 | 23/11/2018 | NA | 3 | 3 | 25 | 0 | 0 | 0 |
| | **2** | 36.081 | -5.626 | 76.781 | 23/11/2018 | NA | 2 | 2 | 15 | 0 | 0 | 0 |
| | **4** | 36.151 | -5.705 | 70.643 | 23/11/2018 | 07/12/2018 | 2 | 2 | 20 | 2 | 2 | 7 |
| | **1** | 36.800 | -5.393 | 10.209 | 23/11/2018 | 11/12/2018 | 30 | 10 | 255 | 22 | 10 | 45 |
| | **5** | 37.259 | -6.097 | 76.421 | 23/11/2018 | 12/12/2018 | 3 | 3 | 30 | 2 | 2 | 7 |
| | **8** | 37.702 | -5.832 | 108.106 | 23/11/2018 | 12/12/2018 | 3 | 3 | 30 | 1 | 1 | 1 |
| | **9** | 37.809 | -6.429 | 142.189 | 23/11/2018 | 12/12/2018 | 2 | 2 | 20 | 2 | 2 | 2 |
| | **10** | 37.882 | -6.618 | 158.788 | 23/11/2018 | 07/02/2019 | 2 | 2 | 20 | 1 | 1 | 1 |

| | | | | | | | | | | | | |
|---|---|---|---|---|---|---|---|---|---|---|---|---|
| | **6** | 37.936 | -5.711 | 131.255 | 23/11/2018 | 18/12/2018 | 30 | 10 | 255 | 17 | 9 | 27 |
| | **CGa1** | 38.217 | 13.322 | 1669.251 | 23/11/2018 | 12/12/2018 | 4 | 3 | 35 | 1 | 1 | 2 |
| | **Ljlb** | 38.221 | -2.454 | 313.956 | 23/11/2018 | NA | 2 | 2 | 20 | 0 | 0 | 0 |
| | **7** | 38.253 | -4.317 | 195.396 | 23/11/2018 | NA | 3 | 3 | 30 | 0 | 0 | 0 |
| | **11** | 38.331 | -3.581 | 242.707 | 23/11/2018 | 12/12/2018 | 3 | 3 | 30 | 1 | 1 | 1 |
| | **19** | 42.171 | -8.684 | 659.678 | 23/11/2018 | 13/12/2018 | 30 | 10 | 270 | 29 | 10 | 159 |
| | **12** | 42.310 | 3.152 | 964.486 | 23/11/2018 | 12/12/2018 | 10 | 9 | 50 | 3 | 3 | 7 |
| | **13** | 43.029 | -3.240 | 721.722 | 23/11/2018 | 11/12/2018 | 5 | 5 | 40 | 5 | 5 | 10 |
| | **Lla** | 43.407 | -4.688 | 740.752 | 23/11/2018 | 08/12/2018 | 4 | 4 | 40 | 4 | 4 | 28 |
| | **15** | 43.463 | -3.653 | 760.076 | 23/11/2018 | 07/12/2018 | 4 | 4 | 40 | 4 | 4 | 30 |
| | **Vil** | 45.094 | -1.050 | 997.500 | 23/11/2018 | 09/12/2018 | 30 | 10 | 223 | 29 | 10 | 113 |
| | **Bro** | 47.167 | -0.207 | 1235.061 | 23/11/2018 | 07/12/2018 | 3 | 3 | 25 | 1 | 1 | 6 |
| | **Roc** | 47.387 | -0.526 | 1248.831 | 23/11/2018 | 10/12/2018 | 3 | 3 | 30 | 3 | 3 | 10 |
| | **Saf** | 47.496 | -1.593 | 1234.486 | 23/11/2018 | 07/12/2018 | 3 | 3 | 30 | 3 | 3 | 20 |
| | **Tal** | 47.700 | -3.455 | 1226.125 | 23/11/2018 | 09/12/2018 | 4 | 4 | 35 | 4 | 4 | 17 |
| | **Mat** | 48.357 | 4.459 | 1522.429 | 23/11/2018 | 08/12/2018 | 30 | 10 | 271 | 30 | 10 | 228 |
| | **BH** | 50.400 | -3.493 | 1523.423 | 23/11/2018 | 09/12/2018 | 3 | 2 | 15 | 2 | 1 | 7 |
| | **IOW** | 50.682 | -1.075 | 1586.039 | 23/11/2018 | 09/12/2018 | 30 | 11 | 175 | 30 | 11 | 113 |
| | **Lil** | 51.202 | -3.187 | 1614.539 | 23/11/2018 | 08/12/2018 | 4 | 2 | 21 | 4 | 2 | 19 |
| | **CR** | 51.278 | -2.795 | 1626.761 | 23/11/2018 | 08/12/2018 | 5 | 3 | 25 | 3 | 2 | 11 |
| | **Man** | 53.137 | -1.144 | 1850.739 | 23/11/2018 | 09/12/2018 | 6 | 6 | 40 | 6 | 6 | 22 |
| | **Tym** | 53.303 | -3.553 | 1844.101 | 23/11/2018 | 09/12/2018 | 7 | 7 | 49 | 7 | 7 | 31 |
| | **Sut** | 53.353 | -0.959 | 1876.825 | 23/11/2018 | 09/12/2018 | 30 | 10 | 225 | 26 | 9 | 128 |
| | **TOTAL** | | | | | | 300 | 154 | 2389 | 242 | 124 | 1052 |
| **Portsmouth** | **3** | 36.036 | -5.556 | 1678.535 | 04/03/2019 | 25/03/2019 | 3 | 3 | 25 | 1 | 1 | 2 |

| | | | | | | | | | | | | |
|---|---|---|---|---|---|---|---|---|---|---|---|---|
| | **2** | 36.081 | -5.626 | 1674.877 | 04/03/2019 | 25/03/2019 | 2 | 2 | 15 | 1 | 1 | 2 |
| | **4** | 36.151 | -5.705 | 1668.659 | 04/03/2019 | 21/03/2019 | 2 | 2 | 20 | 2 | 2 | 4 |
| | **1** | 36.800 | -5.393 | 1592.507 | 04/03/2019 | 24/03/2019 | 30 | 10 | 255 | 26 | 10 | 67 |
| | **5** | 37.259 | -6.097 | 1555.981 | 04/03/2019 | 25/03/2019 | 3 | 3 | 30 | 2 | 2 | 8 |
| | **8** | 37.702 | -5.832 | 1502.737 | 04/03/2019 | 25/03/2019 | 3 | 3 | 30 | 2 | 2 | 11 |
| | **9** | 37.809 | -6.429 | 1503.764 | 04/03/2019 | 25/03/2019 | 2 | 2 | 20 | 1 | 1 | 4 |
| | **10** | 37.882 | -6.618 | 1500.137 | 04/03/2019 | 08/04/2019 | 2 | 2 | 20 | 1 | 1 | 1 |
| | **6** | 37.936 | -5.711 | 1475.092 | 04/03/2019 | 25/03/2019 | 30 | 10 | 255 | 19 | 9 | 29 |
| | **CGa1** | 38.217 | 13.322 | 1801.490 | 04/03/2019 | 25/03/2019 | 4 | 3 | 35 | 2 | 2 | 5 |
| | **Ljlb** | 38.221 | -2.454 | 1401.695 | 04/03/2019 | 25/03/2019 | 2 | 2 | 20 | 2 | 2 | 11 |
| | **7** | 38.253 | -4.317 | 1416.958 | 04/03/2019 | 25/03/2019 | 3 | 3 | 30 | 2 | 2 | 2 |
| | **11** | 38.331 | -3.581 | 1399.155 | 04/03/2019 | 25/03/2019 | 3 | 3 | 30 | 1 | 1 | 2 |
| | **19** | 42.171 | -8.684 | 1120.710 | 04/03/2019 | 25/03/2019 | 30 | 10 | 270 | 27 | 10 | 112 |
| | **12** | 42.310 | 3.152 | 997.814 | 04/03/2019 | 28/03/2019 | 10 | 9 | 50 | 4 | 4 | 4 |
| | **13** | 43.029 | -3.240 | 878.883 | 04/03/2019 | 25/03/2019 | 5 | 5 | 40 | 3 | 3 | 6 |
| | **Lla** | 43.407 | -4.688 | 865.400 | 04/03/2019 | 25/03/2019 | 4 | 4 | 40 | 4 | 4 | 20 |
| | **15** | 43.463 | -3.653 | 838.103 | 04/03/2019 | 25/03/2019 | 4 | 4 | 40 | 4 | 4 | 25 |
| | **Vil** | 45.094 | -1.050 | 634.268 | 04/03/2019 | 26/03/2019 | 30 | 10 | 222 | 27 | 10 | 91 |
| | **Bro** | 47.167 | -0.207 | 409.043 | 04/03/2019 | 25/03/2019 | 3 | 3 | 25 | 2 | 2 | 8 |
| | **Roc** | 47.387 | -0.526 | 381.656 | 04/03/2019 | 25/03/2019 | 3 | 3 | 30 | 2 | 2 | 3 |
| | **Saf** | 47.496 | -1.593 | 369.008 | 04/03/2019 | 22/03/2019 | 3 | 3 | 30 | 3 | 3 | 17 |
| | **Tal** | 47.700 | -3.455 | 384.918 | 04/03/2019 | 25/03/2019 | 4 | 4 | 35 | 4 | 4 | 20 |
| | **Mat** | 48.357 | 4.459 | 484.766 | 04/03/2019 | 24/03/2019 | 30 | 10 | 270 | 30 | 10 | 137 |
| | **BH** | 50.400 | -3.493 | 175.237 | 04/03/2019 | 25/03/2019 | 3 | 2 | 15 | 2 | 1 | 5 |
| | **IOW** | 50.682 | -1.075 | 13.049 | 04/03/2019 | 24/03/2019 | 30 | 11 | 150 | 24 | 11 | 65 |

| | | | | | | | | | | | | |
|---|---|---|---|---|---|---|---|---|---|---|---|---|
| | **Lil** | 51.202 | -3.187 | 153.373 | 04/03/2019 | 25/03/2019 | 4 | 2 | 20 | 2 | 2 | 5 |
| | **CR** | 51.278 | -2.795 | 130.466 | 04/03/2019 | 22/03/2019 | 5 | 3 | 25 | 3 | 2 | 14 |
| | **Man** | 53.137 | -1.144 | 260.280 | 04/03/2019 | 25/03/2019 | 6 | 6 | 40 | 4 | 4 | 11 |
| | **Tym** | 53.303 | -3.553 | 325.619 | 04/03/2019 | 25/03/2019 | 7 | 7 | 49 | 3 | 3 | 12 |
| | **Sut** | 53.353 | -0.959 | 284.413 | 04/03/2019 | 24/03/2019 | 30 | 10 | 225 | 25 | 8 | 106 |
| | **TOTAL** | | | | | | 300 | 154 | 2361 | 235 | 123 | 809 |
| **Durham** | **3** | 36.036 | -5.556 | 2103.551 | 26/03/2019 | 29/04/2019 | 3 | 3 | 25 | 1 | 1 | 1 |
| | **2** | 36.081 | -5.626 | 2099.428 | 26/03/2019 | 29/04/2019 | 2 | 2 | 15 | 1 | 1 | 2 |
| | **4** | 36.151 | -5.705 | 2092.648 | 26/03/2019 | 22/04/2019 | 2 | 2 | 20 | 1 | 1 | 2 |
| | **1** | 36.800 | -5.393 | 2017.599 | 26/03/2019 | 04/05/2019 | 30 | 10 | 255 | 15 | 8 | 20 |
| | **5** | 37.259 | -6.097 | 1975.803 | 26/03/2019 | 29/04/2019 | 3 | 3 | 30 | 2 | 2 | 2 |
| | **8** | 37.702 | -5.832 | 1923.675 | 26/03/2019 | NA | 3 | 3 | 30 | 0 | 0 | 0 |
| | **9** | 37.809 | -6.429 | 1920.130 | 26/03/2019 | 29/04/2019 | 2 | 2 | 20 | 1 | 1 | 1 |
| | **10** | 37.882 | -6.618 | 1914.878 | 26/03/2019 | 29/04/2019 | 2 | 2 | 20 | 2 | 2 | 2 |
| | **6** | 37.936 | -5.711 | 1896.492 | 26/03/2019 | 27/04/2019 | 30 | 10 | 255 | 14 | 7 | 22 |
| | **CGa1** | 38.217 | 13.322 | 2155.785 | 26/03/2019 | 16/05/2019 | 4 | 3 | 35 | 2 | 2 | 5 |
| | **Ljlb** | 38.221 | -2.454 | 1839.928 | 26/03/2019 | 29/04/2019 | 2 | 2 | 20 | 2 | 2 | 6 |
| | **7** | 38.253 | -4.317 | 1846.812 | 26/03/2019 | 22/04/2019 | 3 | 3 | 30 | 1 | 1 | 1 |
| | **11** | 38.331 | -3.581 | 1832.794 | 26/03/2019 | 22/04/2019 | 3 | 3 | 30 | 1 | 1 | 1 |
| | **19** | 42.171 | -8.684 | 1493.716 | 26/03/2019 | 27/04/2019 | 30 | 10 | 270 | 29 | 10 | 113 |
| | **12** | 42.310 | 3.152 | 1427.173 | 26/03/2019 | 29/04/2019 | 10 | 9 | 50 | 2 | 2 | 4 |
| | **13** | 43.029 | -3.240 | 1310.424 | 26/03/2019 | 27/04/2019 | 5 | 5 | 40 | 4 | 4 | 9 |
| | **Lla** | 43.407 | -4.688 | 1282.759 | 26/03/2019 | 22/04/2019 | 4 | 4 | 40 | 4 | 4 | 30 |
| | **15** | 43.463 | -3.653 | 1265.610 | 26/03/2019 | 22/04/2019 | 4 | 4 | 40 | 4 | 4 | 33 |
| | **Vil** | 45.094 | -1.050 | 1076.032 | 26/03/2019 | 23/04/2019 | 30 | 10 | 222 | 29 | 10 | 144 |

| | | | | | | | | | | | |
|---|---|---|---|---|---|---|---|---|---|---|---|
| | **Bro** | 47.167 | -0.207 | 850.348 | 26/03/2019 | 02/05/2019 | 3 | 3 | 25 | 2 | 2 | 9 |
| | **Roc** | 47.387 | -0.526 | 823.752 | 26/03/2019 | 26/04/2019 | 3 | 3 | 30 | 3 | 3 | 13 |
| | **Saf** | 47.496 | -1.593 | 808.363 | 26/03/2019 | 22/04/2019 | 3 | 3 | 30 | 3 | 3 | 26 |
| | **Tal** | 47.700 | -3.455 | 796.543 | 26/03/2019 | 24/04/2019 | 4 | 4 | 35 | 3 | 3 | 18 |
| | **Mat** | 48.357 | 4.459 | 825.702 | 26/03/2019 | 22/04/2019 | 30 | 10 | 272 | 30 | 10 | 220 |
| | **BH** | 50.400 | -3.493 | 502.431 | 26/03/2019 | 24/04/2019 | 3 | 2 | 15 | 3 | 2 | 11 |
| | **IOW** | 50.682 | -1.075 | 455.324 | 26/03/2019 | 25/04/2019 | 30 | 11 | 215 | 29 | 11 | 147 |
| | **Lil** | 51.202 | -3.187 | 410.741 | 26/03/2019 | 22/04/2019 | 4 | 2 | 20 | 3 | 2 | 12 |
| | **CR** | 51.278 | -2.795 | 396.292 | 26/03/2019 | 24/04/2019 | 5 | 3 | 25 | 3 | 2 | 13 |
| | **Man** | 53.137 | -1.144 | 183.063 | 26/03/2019 | 27/04/2019 | 6 | 6 | 40 | 5 | 5 | 22 |
| | **Tym** | 53.303 | -3.553 | 207.791 | 26/03/2019 | 29/04/2019 | 7 | 7 | 49 | 7 | 7 | 28 |
| | **Sut** | 53.353 | -0.959 | 161.963 | 26/03/2019 | 23/04/2019 | 30 | 10 | 226 | 24 | 7 | 138 |
| | **TOTAL** | | | | | | 300 | 154 | 2429 | 230 | 120 | 1055 |
| **OVERALL** | | | | | | | | | | 707 | 367 | 2916 |

### Part 2

| Transplant Site | Population | Survived Pots (Spring) | Survived Families (Spring) | Flowered Pots | Flowered Families | Days to Flowering | Fruited Pots | Fruited Families | Total Fruits | Survived Pots (Autumn) | Survived Families (Autumn) |
|---|---|---|---|---|---|---|---|---|---|---|---|
| **El Bosque** | **3** | 0 | 0 | 0 | 0 | NA | NA | NA | NA | 0 | 0 |
| | **2** | 0 | 0 | 0 | 0 | NA | NA | NA | NA | 0 | 0 |
| | **4** | 2 | 2 | 1 | 1 | 181 | NA | NA | NA | 0 | 0 |
| | **1** | 19 | 9 | 1 | 1 | 169 | NA | NA | NA | 0 | 0 |
| | **5** | 2 | 2 | 0 | 0 | NA | NA | NA | NA | 0 | 0 |
| | **8** | 0 | 0 | 0 | 0 | NA | NA | NA | NA | 0 | 0 |
| | **9** | 2 | 2 | 0 | 0 | NA | NA | NA | NA | 0 | 0 |
| | **10** | 1 | 1 | 0 | 0 | NA | NA | NA | NA | 0 | 0 |

| | | | | | | | | | | | |
|---|---|---|---|---|---|---|---|---|---|---|---|
| | **6** | 13 | 7 | 0 | 0 | NA | NA | NA | NA | 0 | 0 |
| | **CGa1** | 1 | 1 | 0 | 0 | NA | NA | NA | NA | 0 | 0 |
| | **Ljlb** | 0 | 0 | 0 | 0 | NA | NA | NA | NA | 0 | 0 |
| | **7** | 0 | 0 | 0 | 0 | NA | NA | NA | NA | 0 | 0 |
| | **11** | 1 | 1 | 0 | 0 | NA | NA | NA | NA | 0 | 0 |
| | **19** | 27 | 10 | 0 | 0 | NA | NA | NA | NA | 0 | 0 |
| | **12** | 3 | 3 | 0 | 0 | NA | NA | NA | NA | 0 | 0 |
| | **13** | 5 | 5 | 0 | 0 | NA | NA | NA | NA | 0 | 0 |
| | **Lla** | 4 | 4 | 1 | 1 | 171 | NA | NA | NA | 0 | 0 |
| | **15** | 4 | 4 | 0 | 0 | NA | NA | NA | NA | 0 | 0 |
| | **Vil** | 27 | 10 | 5 | 5 | 165 | NA | NA | NA | 0 | 0 |
| | **Bro** | 1 | 1 | 0 | 0 | NA | NA | NA | NA | 0 | 0 |
| | **Roc** | 3 | 3 | 0 | 0 | NA | NA | NA | NA | 0 | 0 |
| | **Saf** | 3 | 3 | 0 | 0 | NA | NA | NA | NA | 0 | 0 |
| | **Tal** | 4 | 4 | 0 | 0 | NA | NA | NA | NA | 0 | 0 |
| | **Mat** | 29 | 10 | 8 | 5 | 167 | NA | NA | NA | 0 | 0 |
| | **BH** | 2 | 1 | 0 | 0 | NA | NA | NA | NA | 0 | 0 |
| | **IOW** | 28 | 11 | 0 | 0 | NA | NA | NA | NA | 0 | 0 |
| | **Lil** | 4 | 2 | 1 | 1 | 168 | NA | NA | NA | 0 | 0 |
| | **CR** | 3 | 2 | 0 | 0 | NA | NA | NA | NA | 0 | 0 |
| | **Man** | 6 | 6 | 0 | 0 | NA | NA | NA | NA | 0 | 0 |
| | **Tym** | 7 | 7 | 0 | 0 | NA | NA | NA | NA | 0 | 0 |
| | **Sut** | 23 | 7 | 0 | 0 | NA | NA | NA | NA | 0 | 0 |
| | **TOTAL** | 224 | 118 | 17 | 14 | 170 | NA | NA | NA | 0 | 0 |
| **Portsmouth** | **3** | 1 | 1 | 1 | 1 | 102 | 1 | 1 | 11 | 1 | 1 |
| | **2** | 1 | 1 | 1 | 1 | 99 | 1 | 1 | 4 | 0 | 0 |
| | **4** | 1 | 1 | 1 | 1 | 102 | 1 | 1 | 27 | 1 | 1 |
| | **1** | 20 | 10 | 20 | 10 | 101 | 18 | 9 | 10 | 18 | 9 |
| | **5** | 2 | 2 | 2 | 2 | 99 | 2 | 2 | 12 | 2 | 2 |

| | | | | | | | | | | | |
|---|---|---|---|---|---|---|---|---|---|---|---|
| | **8** | 2 | 2 | 2 | 2 | 99 | 2 | 2 | 15 | 1 | 1 |
| | **9** | 1 | 1 | 1 | 1 | 99 | 1 | 1 | 4 | 1 | 1 |
| | **10** | 1 | 1 | 1 | 1 | 99 | 1 | 1 | 24 | 1 | 1 |
| | **6** | 15 | 8 | 15 | 9 | 103 | 17 | 9 | 23 | 11 | 8 |
| | **CGa1** | 2 | 2 | 2 | 2 | 99 | 2 | 2 | 11 | 2 | 2 |
| | **Ljlb** | 2 | 2 | 2 | 2 | 118 | 2 | 2 | 5 | 1 | 1 |
| | **7** | 2 | 2 | 2 | 2 | 104 | 2 | 2 | 13 | 2 | 2 |
| | **11** | 1 | 2 | 1 | 1 | 99 | 1 | 1 | 9 | NA | 2 |
| | **19** | 22 | 10 | 22 | 10 | 128 | 16 | 9 | 8 | 19 | 10 |
| | **12** | 3 | 3 | 3 | 3 | 117 | 3 | 3 | 8 | 3 | 3 |
| | **13** | 2 | 2 | 2 | 2 | 115 | 2 | 2 | 10 | 1 | 1 |
| | **Lla** | 4 | 4 | 4 | 4 | 114 | 4 | 4 | 14 | 2 | 2 |
| | **15** | 3 | 4 | 3 | 3 | 122 | 2 | 2 | 11 | NA | 4 |
| | **Vil** | 26 | 9 | 26 | 9 | 114 | 25 | 9 | 13 | 25 | 9 |
| | **Bro** | 2 | 2 | 1 | 1 | 162 | 1 | 1 | 2 | 2 | 2 |
| | **Roc** | 2 | 2 | 2 | 2 | 108 | 2 | 2 | 25 | 2 | 2 |
| | **Saf** | 3 | 3 | 3 | 3 | 128 | 3 | 3 | 5 | 3 | 3 |
| | **Tal** | 4 | 4 | 4 | 4 | 114 | 3 | 3 | 4 | 4 | 4 |
| | **Mat** | 27 | 10 | 27 | 10 | 112 | 26 | 10 | 18 | 24 | 10 |
| | **BH** | 1 | 1 | 1 | 1 | 115 | 1 | 1 | 80 | 1 | 1 |
| | **IOW** | 22 | 12 | 21 | 10 | 117 | 21 | 10 | 11 | NA | 12 |
| | **Lil** | 2 | 2 | 2 | 2 | 115 | 2 | 2 | 17 | 2 | 2 |
| | **CR** | 3 | 2 | 3 | 2 | 109 | 3 | 2 | 15 | 3 | 2 |
| | **Man** | 3 | 3 | 3 | 3 | 112 | 3 | 3 | 19 | 3 | 3 |
| | **Tym** | 2 | 3 | 2 | 2 | 128 | 2 | 2 | 5 | NA | 3 |
| | **Sut** | 21 | 8 | 20 | 8 | 115 | 19 | 8 | 9 | 20 | 8 |
| | **TOTAL** | 203 | 119 | 200 | 114 | 112 | 189 | 110 | 442 | 155 | 112 |
| **Durham** | **3** | 1 | 1 | 1 | 1 | 154 | NA | NA | NA | 1 | 1 |
| | **2** | 0 | 0 | 0 | 0 | NA | NA | NA | NA | 0 | 0 |

| | | | | | | | | | | |
|---|---|---|---|---|---|---|---|---|---|---|
| **4** | 1 | 1 | 1 | 1 | 174 | NA | NA | NA | 1 | 1 |
| **1** | 9 | 5 | 0 | 0 | NA | NA | NA | NA | 4 | 3 |
| **5** | 2 | 2 | 0 | 0 | NA | NA | NA | NA | 1 | 1 |
| **8** | 0 | 0 | 0 | 0 | NA | NA | NA | NA | 0 | 0 |
| **9** | 0 | 0 | 0 | 0 | NA | NA | NA | NA | 0 | 0 |
| **10** | 2 | 2 | 0 | 0 | NA | NA | NA | NA | 0 | 0 |
| **6** | 9 | 7 | 2 | 1 | 139 | NA | NA | NA | NA | 4 |
| **CGa1** | 2 | 2 | 0 | 0 | NA | NA | NA | NA | 0 | 0 |
| **Ljlb** | 2 | 2 | 0 | 0 | NA | NA | NA | NA | 1 | 1 |
| **7** | 0 | 0 | 0 | 0 | NA | NA | NA | NA | 0 | 0 |
| **11** | 1 | 1 | 0 | 0 | NA | NA | NA | NA | 0 | 0 |
| **19** | 26 | 10 | 0 | 0 | NA | NA | NA | NA | 22 | 10 |
| **12** | 1 | 1 | 0 | 0 | NA | NA | NA | NA | 1 | 1 |
| **13** | 4 | 4 | 0 | 0 | NA | NA | NA | NA | 1 | 1 |
| **Lla** | 4 | 4 | 0 | 0 | NA | NA | NA | NA | 3 | 3 |
| **15** | 4 | 4 | 0 | 0 | NA | NA | NA | NA | 2 | 2 |
| **Vil** | 28 | 10 | 2 | 1 | 136 | NA | NA | NA | 19 | 9 |
| **Bro** | 1 | 1 | 0 | 0 | NA | NA | NA | NA | 1 | 1 |
| **Roc** | 3 | 3 | 0 | 0 | NA | NA | NA | NA | 3 | 3 |
| **Saf** | 3 | 3 | 1 | 1 | 168 | NA | NA | NA | 3 | 3 |
| **Tal** | 3 | 3 | 0 | 0 | NA | NA | NA | NA | 3 | 3 |
| **Mat** | 27 | 10 | 1 | 1 | 155 | NA | NA | NA | 21 | 10 |
| **BH** | 3 | 2 | 0 | 0 | NA | NA | NA | NA | 1 | 1 |
| **IOW** | 27 | 11 | 2 | 2 | 142 | NA | NA | NA | 22 | 10 |
| **Lil** | 2 | 2 | 0 | 0 | NA | NA | NA | NA | 2 | 2 |
| **CR** | 3 | 2 | 0 | 0 | NA | NA | NA | NA | 2 | 1 |
| **Man** | 4 | 4 | 1 | 1 | 171 | NA | NA | NA | 4 | 4 |
| **Tym** | 6 | 6 | 0 | 0 | NA | NA | NA | NA | 5 | 5 |
| **Sut** | 23 | 6 | 1 | 1 | 154 | NA | NA | NA | 20 | 6 |

| | | | | | | | | | | |
|---|---|---|---|---|---|---|---|---|---|---|
| | **TOTAL** | 201 | 109 | 12 | 10 | 155 | NA | NA | NA | 143 | 86 |
| **OVERALL** | | 628 | 346 | 229 | 138 | 146 | NA | NA | NA | 298 | 198 |

Supplementary table 4. Results of the principal component analyses, including the loadings and principal components PC1 to PC3, to summarise the local climate of the populations surveyed along the latitudinal gradient. Positive and negative loadings contributing more than 0.5 to each PC are coloured in blue and in red, respectively.

| Loadings | | | |
|---|---|---|---|
| **ClimatVar_Season** | **PC1** | **PC2** | **PC3** |
| prec_DJF | 0.174 | 0.529 | -0.681 |
| prec_JJA | -0.869 | 0.209 | -0.273 |
| prec_MAM | -0.094 | 0.445 | -0.849 |
| prec_SON | -0.233 | 0.615 | -0.694 |
| srad_DJF | 0.930 | -0.227 | -0.008 |
| srad_JJA | 0.894 | -0.269 | 0.051 |
| srad_MAM | 0.930 | -0.224 | 0.020 |
| srad_SON | 0.940 | -0.200 | -0.032 |
| tavg_DJF | 0.922 | 0.352 | -0.029 |
| tavg_JJA | 0.923 | -0.298 | 0.158 |
| tavg_MAM | 0.991 | 0.035 | 0.024 |
| tavg_SON | 0.992 | 0.088 | 0.046 |
| tmax_DJF | 0.964 | 0.125 | -0.042 |
| tmax_JJA | 0.807 | -0.523 | 0.128 |
| tmax_MAM | 0.947 | -0.253 | 0.003 |
| tmax_SON | 0.974 | -0.167 | 0.025 |
| tmin_DJF | 0.783 | 0.591 | -0.012 |
| tmin_JJA | 0.959 | 0.111 | 0.183 |
| tmin_MAM | 0.913 | 0.387 | 0.042 |
| tmin_SON | 0.911 | 0.384 | 0.068 |
| vapr_DJF | 0.827 | 0.501 | -0.030 |
| vapr_JJA | 0.638 | 0.589 | 0.049 |
| vapr_MAM | 0.810 | 0.523 | -0.018 |
| vapr_SON | 0.805 | 0.532 | -0.002 |
| wind_DJF | -0.545 | 0.684 | 0.434 |
| wind_JJA | -0.377 | 0.625 | 0.544 |
| wind_MAM | -0.453 | 0.690 | 0.489 |
| wind_SON | -0.517 | 0.653 | 0.504 |

| Principal Components | | | |
|---|---|---|---|
| **Population** | **PC1** | **PC2** | **PC3** |
| 3 | 5.367 | 3.009 | 0.541 |
| 2 | 6.383 | 3.188 | -0.130 |
| 4 | 6.533 | 2.432 | -0.611 |
| 1 | 1.846 | -1.332 | -0.190 |
| 5 | 6.354 | -1.009 | 0.370 |
| 8 | 5.655 | -1.622 | 0.537 |
| 9 | 3.087 | -2.111 | 0.259 |
| 10 | 2.429 | -2.026 | 0.143 |
| 6 | 2.619 | -2.285 | 0.324 |
| 7 | 1.175 | -3.868 | 0.203 |
| 11 | 1.613 | -4.139 | 0.844 |
| 19 | -1.551 | 2.828 | -5.173 |
| 12 | 0.120 | 1.792 | 3.345 |
| 14 | -3.398 | -3.411 | -1.836 |
| 13 | -3.830 | -2.266 | -2.315 |
| 15 | -0.135 | 1.706 | -2.983 |
| IOW2 | -5.447 | 2.161 | 1.706 |
| IOW1 | -5.463 | 2.189 | 1.743 |
| Tor | 6.561 | 1.674 | 2.282 |
| CGa1 | 5.845 | 2.724 | 2.514 |
| Lla | -0.093 | 0.232 | -1.601 |
| Vil | -1.449 | 0.359 | -0.876 |
| Bro | -3.291 | -0.895 | 0.205 |
| Roc | -3.057 | -0.802 | 0.147 |
| Saf | -2.820 | -0.663 | -0.468 |
| Tal | -3.097 | 3.614 | 1.631 |
| Mat | -4.574 | -2.090 | -0.497 |
| BH | -5.397 | 3.754 | 2.368 |
| Lil | -5.769 | 1.930 | 0.624 |
| CR | -5.348 | 0.097 | -0.455 |
| Tym | -6.617 | 1.478 | 1.481 |
| Sut | -6.676 | -0.878 | 1.388 |
| Man | -7.323 | -0.454 | 0.932 |
| LJLb1 | 0.054 | -4.368 | 1.209 |
| Dor | -6.279 | 2.961 | 2.316 |

SUPPLEMENTARY FIGURES

Supplementary figure 1. Pearson's coefficients of correlation, degrees of freedom, and statistical significance are reported for the correlation between mean flowering initiation by population scored across Experiment 1, Experiment 2 (no vernalization and vernalization are considered separately), and Experiment 3 (Portsmouth site).

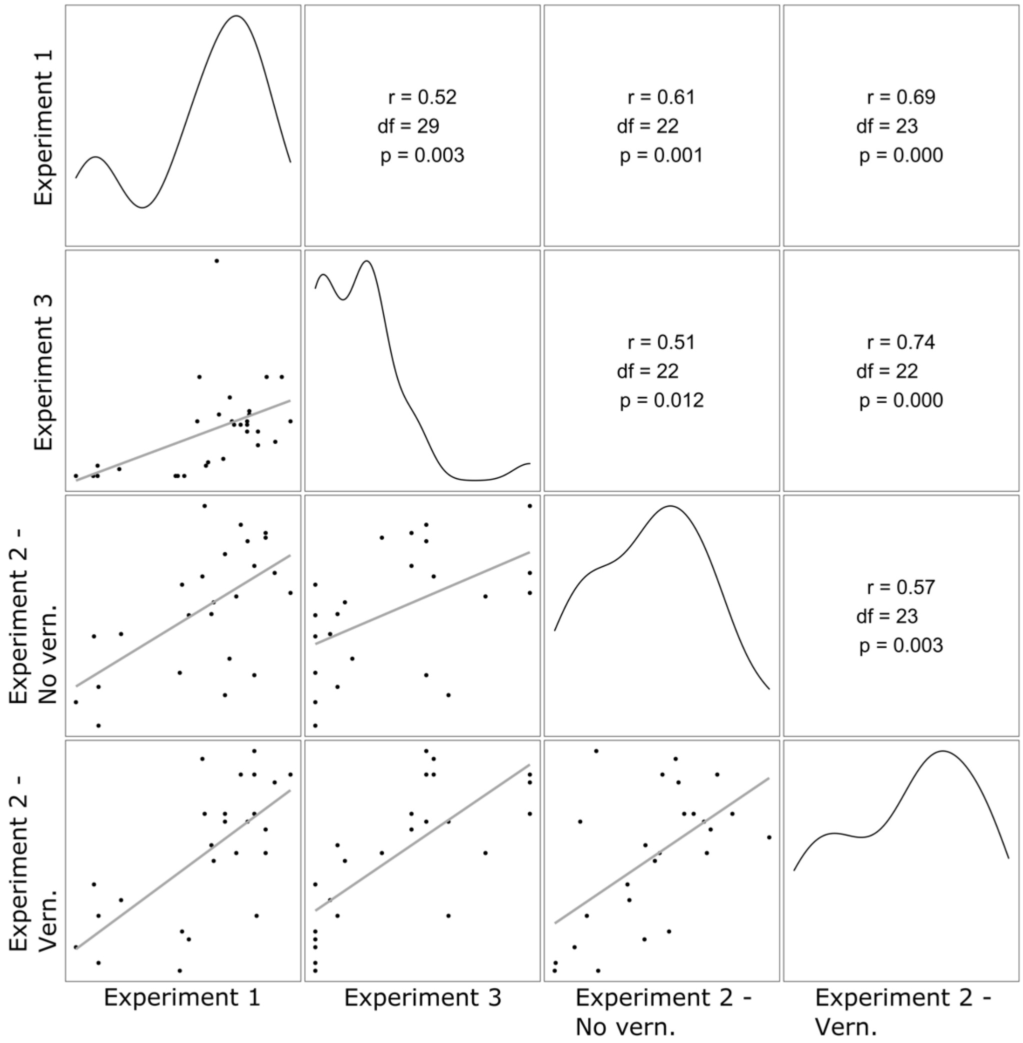

Supplementary figure 2. Evanno's method plot to retrieve optimal number of clusters for *Linum. bienne* based on STRUCTURE output. (A) Mean likelihood for each K value, (B) Rate of change of the likelihood distribution, (C) Absolute values of the second order rate of change of the likelihood distribution, (D) Number of possible clusters, Delta K = 2 indicates the maximum K value. An examination of all parameters shows that there is an agreement with the optimal number of clusters K being 2.

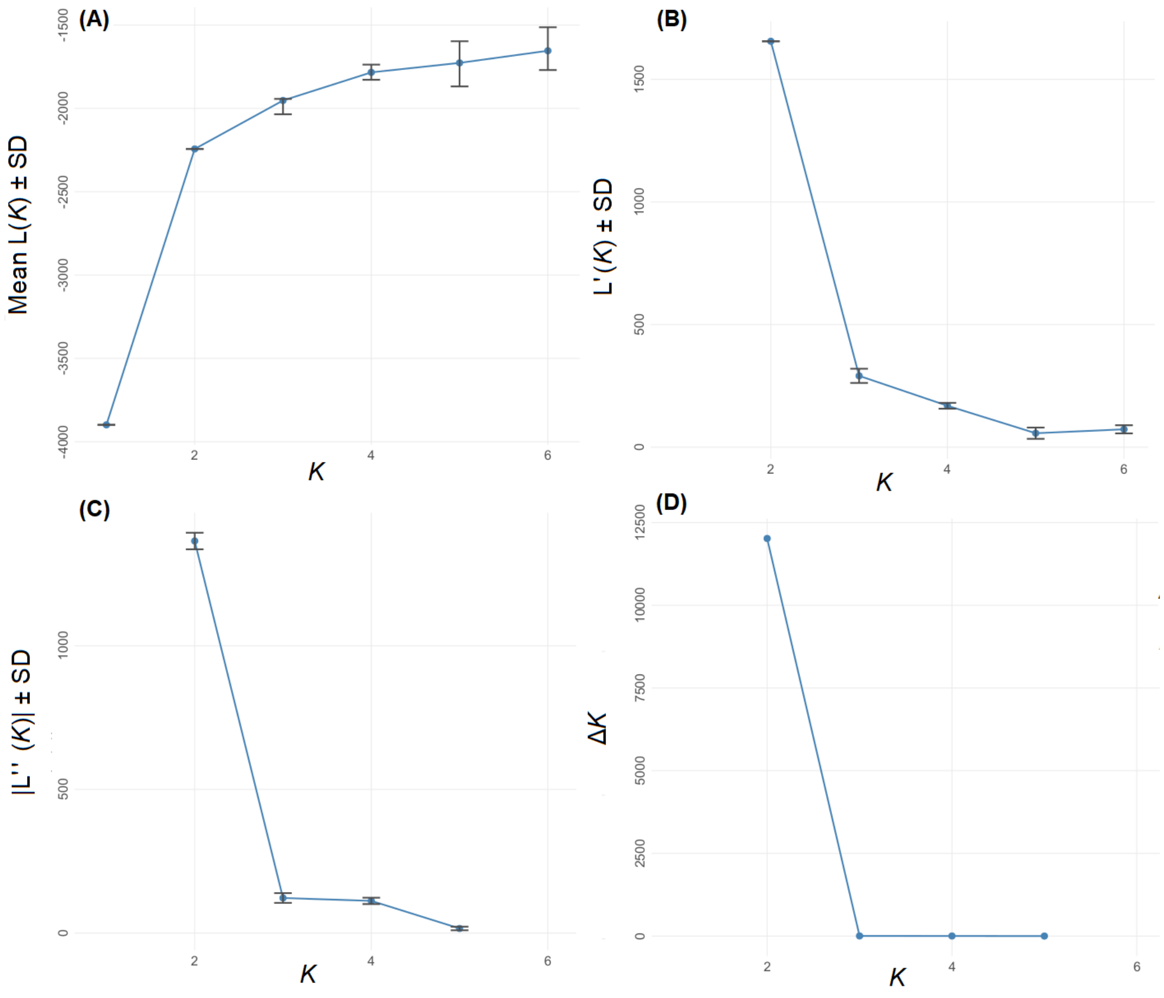
